# C9orf50 is a Targetable Spliceosome Regulator for Cancer Immunotherapy

**DOI:** 10.1101/2025.03.13.642772

**Authors:** Tong Shao, Chuanyang Liu, Jingyu Kuang, Sisi Xie, Ying Qu, Yingying Li, Fangzhou Liu, Lulu Zhang, Ming Li, Tao Hou, Yanhua Qi, Jingyang Wang, Sujuan Zhang, Yu Liu, Jiali Liu, Yanming Hu, Shaowei Zhang, Lingyun Zhu, Jianzhong Shao, Aifu Lin, Guangchuan Wang, Lvyun Zhu

## Abstract

Cancer cells harbor cell-intrinsic programs capable of sculpting immunogenic landscapes, yet molecular brakes that enforce immune evasion remain poorly defined. Here, we identify C9ORF50, a previously uncharacterized intrinsically disordered protein, as a liquid-liquid phase separation-driven regulator of RNA processing and actionable immunotherapy target. Genetic ablation of C9ORF50 induces selective intron retention in spliceosome component transcripts, leading to cytoplasmic accumulation of immunogenic double-stranded RNA (dsRNA). This dsRNA activates type I interferon pathway that in turn orchestrates a chemokine secretory program critical for T cell recruitment. The resultant enhancement of tumor immunogenicity and reprogramming of tumor immune microenvironment drives robust antitumor immunity, effectively converting immunologically “cold” tumors to “hot” phenotypes. Therapeutic targeting of C9ORF50 via RNA interference achieved robust tumor suppression in syngeneic models, suggesting its translational relevance. Notably, C9ORF50 exhibits tumor-predominant expression, with minimal baseline levels detected in healthy tissues or immune cell populations, underscoring its potential safety as a therapeutic target. Our study not only identifies C9ORF50 as a novel LLPS-dependent regulator of RNA splicing but also establishes it as a promising and potentially safe therapeutic target for cancer immunotherapy.

## Introduction

Cancer immunotherapies, such as immune checkpoint blockade (ICB), CAR-T cell therapy, and cancer vaccines, have revolutionized oncology treatments ^1–3^, achieving unprecedented success across various cancer types, particularly in hematological malignancies. However, the durable responses remain limited to subsets of patients ^4^. The effectiveness of immunotherapy is intricately linked to the cancer cell-intrinsic mechanisms, which determines whether the primary tumor is eradicated, metastasizes, or persists as dormant micro-metastases ^5^. A pivotal mechanism in cancer’s ability to evade immune detection and resist immunotherapy involves the regulation of cancer cell-intrinsic mechanisms ^6, 7^. These mechanisms orchestrate cell-intrinsic immune responses, potentially enabling the tumor to sidestep immune surveillance by fostering an immune-privileged tumor microenvironment (TME) ^6, 8^. In this context, T cells and other effective immune cells are often excluded, thereby suppressing a robust anti-tumor immune response ^5, 9, 10^. Therefore, identifying druggable regulators that intrinsically bridge tumor cell biology with immune recognition represents an unmet therapeutic frontier.

Aberrant alternative RNA splicing, a hallmark of cancer, fuels tumorigenesis and therapy resistance while representing a source of diagnostic biomarkers and actionable therapeutic targets ^1111^. Mounting evidences implicate the splicing factors, such as SRSF6, SF3A2 and SF3B1, play pivotal roles in fueling malignant progression ^12–15^, motivating pharmacological targeting of spliceosomal machinery. Clinical-stage small molecules such as SF3B1 inhibitors (E7107 and H3B-8800) disrupt branch site recognition to selectively modulate survival-associated transcripts ^16–18^, whereas herboxidiene exerts antitumor effects by blocking tri-snRNP recruitment, thereby dysregulating invasion and immune-related pathways ^19^. Beyond direct inhibition, degraders like E7820 hijack the CUL4-DCAF15 ubiquitin ligase complex to eliminate RBM39, demonstrating efficacy in metastatic malignancies ^20^. Notably, emerging studies reveal synergistic potential between splicing modulation and immunotherapy: RBM39 degradation or arginine methyltransferase (RMT) inhibition enhances tumor antigen presentation, a mechanism proposed to counteract immune checkpoint blockade (ICB) resistance ^21^. Despite these advances, clinical translation remains challenging due to the inherent complexity of spliceosome networks, tumor heterogeneity and toxicity concerns.

Intrinsically disordered proteins (IDPs), which lack stable tertiary structures, orchestrate RNA splicing through liquid-liquid phase separation (LLPS), dynamically organizing spliceosome components into membraneless condensates ^22–24^. While IDP-driven LLPS enables spatiotemporal control of spliceosome assembly and transcript processing, its functional interplay with antitumor immunity remains unexplored. In sarcoma, EWSR1-FLI1 exploits phase separation to hijack transcriptional programs, generating oncogenic chimeric transcripts that drive malignant transformation ^25, 26^. However, whether IDP-mediated splicing directly modulates antitumor immunity remains uncharted. Unraveling this interplay could provide novel insights into immune evasion mechanisms and inform the development of splicing-targeted immunotherapies.

In this study, we identify C9ORF50, a previously uncharacterized gene, as a potential targetable oncogene for cancer therapy through genome-wide *in vivo* CRISPR screen in immunocompetent syngeneic model. Structurally defined as an IDP, C9ORF50 licenses antitumor immunity by mediating LLPS-dependent regulation of RNA splicing. Genetic perturbation of C9ORF50 activates cancer cell-intrinsic innate immune pathways through splicing dysregulation, culminating in the conversion of immunologically “cold” to “hot” tumors via enhanced T cell infiltration and amplified antitumor immunity. Our findings not only establish C9ORF50 as a druggable target that bridges RNA splicing fidelity to immune surveillance but also introduce a novel paradigm for uncovering tumor cell-intrinsic immunomodulators and advancing next-generation splicing-targeted immunotherapies.

## Results

### Genome-scale in vivo CRISPR screen identifies C9ORF50 as a critical driver of cancer progression

To identify key genes involved in cancer progression under immunosurveillance, we performed a genome-wide *in vivo* CRISPR knockout screen in immunocompetent C57BL/6 mice using Cas9 stably expressing MC38 cells transduced with the mBrie library (**Figure 1A**). While MC38-Cas9-mBrie and MC38-Cas9-NTC control tumors (non-targeting control sgRNA-transduced MC38-Cas9 cells) exhibited comparable growth through day 9, mBrie tumors regressed completely by day 20, implicating rejection outcomes of certain gene-knockout tumor cells (p < 0.001; **Figures S1A and S1B**). To identify specific genes responsible for tumor inhibition, we harvested MC38-Cas9-mBrie tumors at day 9 post-inoculation, extracted genomic DNA, and quantified sgRNA abundance through sequencing. Parallel sequencing of pre-injection MC38-Cas9-mBrie cells and the original plasmid library revealed high coverage and strong inter-library correlation (**Figures S1C and S1D**). The consistent sgRNA distribution across replicates and distinct profiles between tumor and cell libraries demonstrated robust screening performance and significant *in vivo* selective pressure (**Figures S1E and S1F**). Using RIGER and MAGeCK algorithms for sgRNA enrichment analysis, we identified 261 and 268 genes, respectively, showing significant negative selection (**Figure S1G**). Convergent analysis yielded 15 high-confidence candidates associated with tumor rejection, including established immunotherapy targets (Alk, Spi1^27, 28^), cytoskeleton regulators (Baiap2, 2610034B18Rik/Arpin), mitochondrial components (Chchd4, Cox5a, Hsd17b10, Slc25a26), histone chaperones (Nap1l4, Spty2d1), RNA processing factors (Parn, Tra2a), and uncharacterized genes (1700001O22Rik/C9ORF50, Vmn1r191) (**Figure S1H**). sgRNA read analysis confirmed marked depletion of all 15 candidates in tumors, surpassing a stringent FDR threshold of 0.2%. A direct comparison of sgRNA reads revealed marked depletion of sgRNAs targeting these 15 genes in tumors, with at least one sgRNA surpassing a stringent false discovery rate (FDR) threshold of 0.2% (**Figures 1B, S1I-S1K**). For validation, we selected eight candidates (C9ORF50, Baiap2, Cox5a, Slc25a26, Spty2d1, Tra2a, Parn, Nap1l4) and generated individual knockout constructs. T7E1 assays confirmed efficient knockout **(Figure S2A)**, and monoclonal knockout lines demonstrated consistent tumor suppression in immunocompetent C57BL/6 mice compared to NTC controls **(Figure S2B)**. As additional validation, knockout of known tumor suppressors SetD2 and Smad4 maintained normal tumor growth, confirming the specificity of our approach **(Figure S2C)**. Collectively, this *in vivo* screen robustly identified high-confidence candidate genes critical for tumor growth under immune pressure.

**Figure 1.**
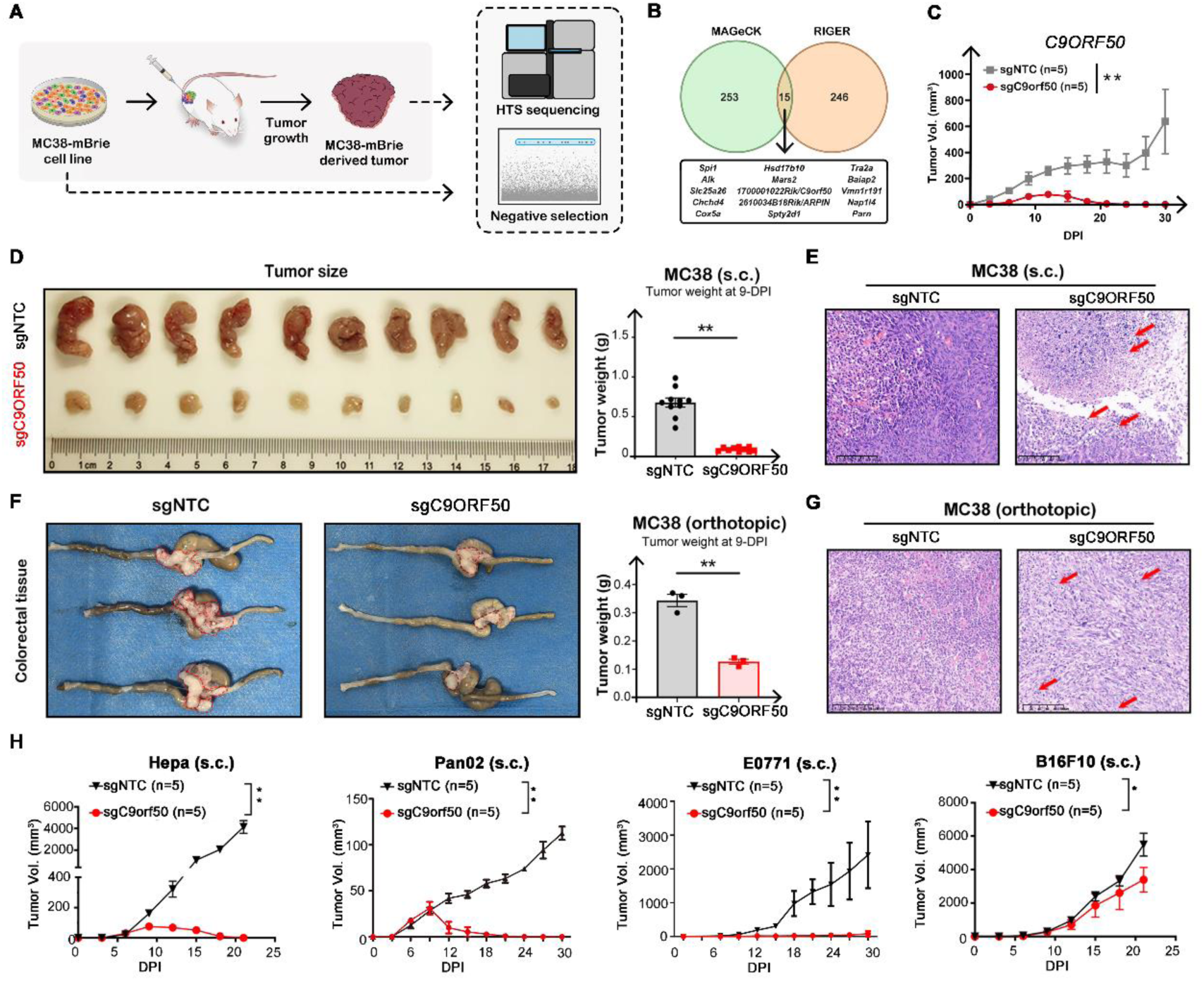
Genome-scale *in vivo* CRISPR screen identifies C9ORF50 as a critical driver of cancer progression. (A) Schematics of the experimental design of CRISPR knockout screen. (B) Venn diagram of the overlap of hits identified using RIGER analysis and MAGeCK analysis, revealing 15 genes significant regardless of criteria. (C) Tumor growth curve of subcutaneously transplanted MC38 cells transduced with C9ORF50-knockout sgRNA (sgC9ORF50, n = 10 mice) or NTC sgRNA (sgNTC, n = 10 mice) in C57BL/6 mice. **p < 0.01 by two-way ANOVA. Data are shown as mean ± SEM. (D) Representative photographs and weights of tumors from heterotopic tumor transplant. **p < 0.01 by Student t-test. Data are shown as mean ± SEM. **p < 0.01 by two-way ANOVA. Data are shown as mean ± SEM. (E) Representative H&E images of tumors from (d). Scale bar, 100 μm. (F) Representative photographs and weights of tumors with orthotopic tumor transplant. Tumors were circled out by dashed lines. **p < 0.01 by T.test. Data are shown as mean ± SEM. (G) Representative H&E images of tumors in (D). Scale bar, 100 μm. (H) Tumor growth curve of subcutaneously transplanted Hepa1-6, Pan02,E0771, and B16F10 cells transduced with C9ORF50-knockout sgRNA (sgC9ORF50, n=5 mice) or NTC sgRNA (n=5 mice) in C57BL/6 mice. **p < 0.01 by twoway ANOVA. Data are shown as mean ± SEM.

Given the limited understanding of the uncharacterized gene C9ORF50 identified in our screen, we sought to explore its role in tumor growth and immunosurveillance. In the heterotopic model, C9ORF50 knockout MC38 cells were transplanted subcutaneously into mice alongside NTC controls. Tumor growth curves demonstrated a significant reduction in tumor progression in the C9ORF50 knockout group (**Figure 1C**). On day 9 post-transplantation, tumors were excised for direct measurement, revealing that C9ORF50-knockout tumors were notably smaller (**Figures 1D, S3A and S3B**). Histological analysis further confirmed a lower cell density and pronounced leukocyte infiltration in C9ORF50-knockout tumors (**Figures 1E and S3C**). Similarly, in orthotopic transplantation models, C9ORF50 knockout significantly reduced tumor growth compared to the NTC controls (**Figures 1F, 1G, and S3D**). These results indicate that C9ORF50 knockout effectively impedes tumor progression *in vivo*.

To further investigate whether C9ORF50 knockout affects tumor growth in various cancer types, we knocked out C9ORF50 in several cell lines, including Hepa1-6 (Hepatoma), Pan02 (pancreatic cancer), E0771 (triple-negative breast cancer), and B16F10 (melanoma), and transplanted these cells into C57BL/6 mice (**Figure S3E**). Consistent with our observations in the MC38 model, C9ORF50 knockout in these cells significantly inhibited tumor growth and resulted in better survival compared to NTC controls (**Figures 1H and S3F**). Moreover, tissue distribution analysis revealed elevated C9ORF50 expression in the stomach, intestine, and brain, in contrast to diminished levels in the heart, spleen, kidney, and liver, implying a tissue-specific response to C9ORF50 ablation, potentially influencing tumor rejection efficacy (**Figure S3G**). Collectively, these findings suggest that C9ORF50 knockout enhances tumor rejection in various cancer types, particularly those of gastrointestinal origin.

### C9ORF50 is a canonical IDP driving LLPS

Structural analysis revealed that human and mouse C9ORF50 harbor a conserved domain of unknown function (DUF4685) but lack resolvable tertiary architecture **(Figures 2A, S4A-B)**. Computational disorder profiling using IUPred2, ANCHOR2, and MetaPredict revealed extensive intrinsically disordered regions (IDRs) **(Figure 2B)**. Biophysical characterization confirmed hallmark IDP signatures: both human and mouse C9ORF50 exhibited low order-promoting residue (OPR) content (human: 27.1%; mouse: 28.6%) and high disorder-promoting residue (DPR) enrichment (human: 62.2%; mouse: 57.5%), values comparable to canonical IDPs like alpha-synuclein, osteopontin, emerin and TFIP11 **(Figure 2C)**. Charge-hydropathy analysis further classified C9ORF50 within the disordered protein regime (human: net charge = 0.0680, hydropathy = 0.4228; mouse: net charge = 0.0800, hydropathy = 0.3987) **(Figures 2B-D)**. These features suggest C9ORF50 is a canonical IDP.

**Figure 2.**
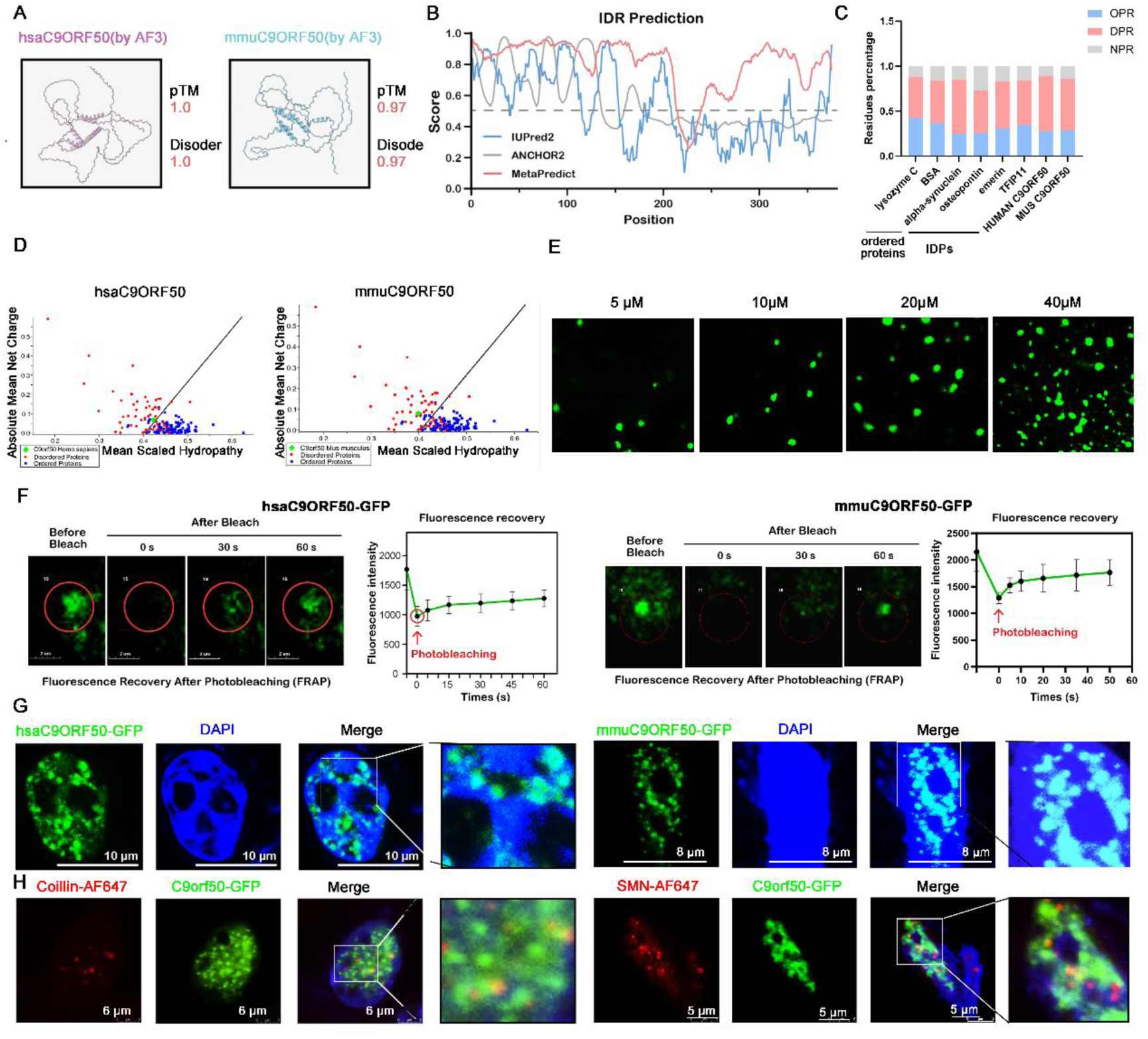
C9ORF50 is a canonical IDP driving LLPS. (A) Three-dimensional (3-D) structures of the C9ORF50 orthologue genes in human (B) IUPred2, ANCHOR2 and MetaPredict analysis the disordered regions (IDRs) in C9ORF50. (C) Occurrence of OPR (order-promoting residues), DPR (disorder-promoting residues) and NPR (non-promoting disorder/order residues) in the sequence of well-ordered proteins (lysozyme C, BSA), known IDPs (Osteopontin, α-Synuclein, Emerin) and C9ORF50. (D) Charge-Hydropathy plot of the hsaC9ORF50 and mmuC9ORF50 using the PONDR algorithm. The solid black line represents the border between ordered and disordered phases. (E) C9ORF50 fused into liquid droplets in a concentration-dependent manner *in vitro*. Scale bar, 5 μm. (E) Subcellular localization of husC9ORF50-GFP and mmuC9ORF50-GFP in MC38 cells. Scale bar, 10 μm. (F) The fluorescence intensity of husC9ORF50-GFP (left) and mmuC9ORF50-GFP (right) droplets recovered after bleaching during FRAP assay. Quantification of fluorescence intensity recovery in the bleached region of C9ORF50 droplets (Right). Time 0 indicates the photobleaching pulse. Scale bar, 1 μm. Error bars, SEM of three independent experiments. (G) Subcellular localization of husC9ORF50-GFP and mmuC9ORF50-GFP in MC38 cells. Scale bar, 10 μm. (H) Representative micrographs of C9ORF50-GFP puncta before and after photobleaching. Scale bar, 2 μm. Quantification of fluorescence intensity recovery in the bleached region of C9ORF50-GFP puncta. Error bars, SEM of six independent experiments.

Given IDRs’ role in LLPS, we reconstituted C9ORF50 phase separation behavior *in vitro*. Purified mouse and human C9ORF50-GFP (5–40 μM) formed spherical condensates that fused into larger structures over time and was concentration-dependent **(Figure 2E)**. FRAP assays in living cells revealed dynamic nuclear puncta of C9ORF50-GFP exhibiting fusion events, with 38.1%(human) and 55.1%(mouse) fluorescence recovery within 60 seconds post-bleaching **(Figure 2F)**. Super-resolution microscopy showed that C9ORF50 was colocalized with coilin-positive Cajal bodies (CBs) and SMN-positive Gemini of Cajal bodies (GEMs), subnuclear hubs governing spliceosome assembly **(Figures 2G and 2H)**. These results suggest that C9ORF50 undergoes LLPS to form dynamic nuclear condensates which may corelate with spliceosome assembly.

### C9ORF50 interacts with spliceosome components and regulates their expression

To comprehensively characterize the intracellular interaction network of C9ORF50, we performed co-immunoprecipitation followed by mass spectrometry analysis (CoIP-MS). Our proteomic profiling revealed significant enrichment of RNA processing and modification factors associated with C9ORF50 **(Figures 3A, 3B, S5B and S5C)**. Subsequent validation through CoIP-Western blot analysis confirmed robust interactions between C9ORF50 and key spliceosome components, including U2SURP and SRSF12 **(Figures 3B and 3C)**. Furthermore, immunofluorescence microscopy demonstrated substantial co-localization of endogenous spliceosome factors, such as U2SURP, SRSF12, SF3B1, and U2AF65, with C9ORF50-GFP puncta within nuclear compartments **(Figures 3D and 3E)**. These findings collectively suggest that C9ORF50 may orchestrate RNA splicing events through the formation of LLPS-mediated nuclear condensates.

**Figure 3.**
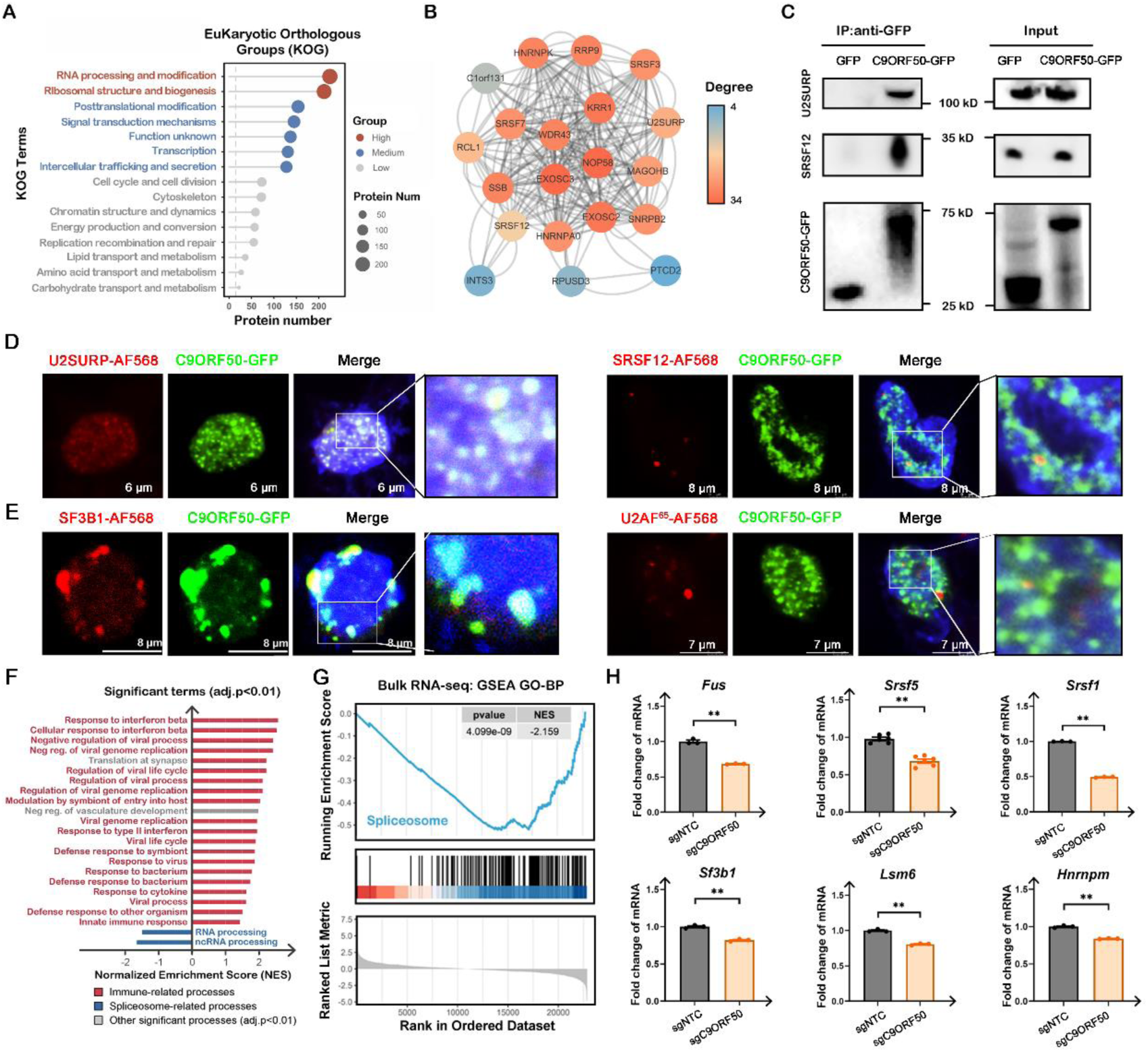
C9ORF50 interacts with spliceosome components and regulates their expression. (A) Clusters of Orthologous Groups (COG)/ Eukaryotic Orthologous Groups (KOG) enrichment analyses the protein set enrichment of interacting with the C9ORF50. (B) The interaction network that were interacted with the C9ORF50. The complex network was constructed by the STRING database, and illustrated with the Cytoscape software. (C) Western blot analysis confirms that both U2SURP and SRSF12 interacted with the C9ORF50. 293T cells were transfected with pCMV-C9ORF50-GFP or pCMV-GFP plasmids. Cell lysates were immunoprecipitated with anti-GFP antibody, followed by immunoblotting analysis with antibodies against U2SURP and SRSF12. SDHA served as loading control. (D) Representative U2SURP, SRSF12 immunofluorescence staining in C9ORF50-GFP positive MC38 cells. Scale bar, 8 μm. Images representative of 3 experiments. (E) Representative SF3B1, U2AF^65^ immunofluorescence staining in C9ORF50-GFP positive MC38 cells. Scale bar, 8 μm. Images representative of 3 experiments. (F) Hierarchical clustering of genes significantly changed in C9ORF50 knockout MC38 cells compared with that in NTC MC38 cells by RNA-seq. (G) GSEA analyzed the spliceosome pathway genes set in C9ORF50 knockout MC38 cells compared with that in NTC MC38 cells. (H) Expression of Fus, Srsf5, Srsf1, Sf3b1, Lsm6 and Hnrnpm in C9ORF50 knockout versus NTC MC38 cells were measured by qRT-PCR. *p < 0.05, **p < 0.01 by Student’s t test. Data are shown as mean ± SEM.

To systematically assess the functional consequences of C9ORF50 deficiency on RNA splicing, we performed comprehensive transcriptomic profiling comparing C9ORF50 knockout and NTC MC38 cells. Our analysis identified 841 differentially expressed genes (DEGs), comprising 510 downregulated and 331 upregulated transcripts in C9ORF50-deficient cells **(Figure S6)**. Notably, gene ontology (GO) analysis revealed a striking dichotomy in functional enrichment: upregulated genes were predominantly associated with immune-related processes, while downregulated genes showed significant enrichment in mRNA splicing pathways **(Figures 3F and S6)**. Further pathway analysis using the Kyoto Encyclopedia of Genes and Genomes (KEGG) database confirmed the significant downregulation of genes involved in spliceosome-related pathways **(Figures 3G and S7)**. To validate these findings, we performed quantitative PCR (qPCR) analysis of key splicing factors, including Fus, Srsf5, Srsf1, Sf3b1, Lsm6, and Hnrnpm, which consistently demonstrated reduced expression levels in C9ORF50 knockout cells **(Figure 3H)**. Taken together, these results provide compelling evidence that C9ORF50 serves as a critical regulator of spliceosome function through directly interacting with spliceosome and maintaining the expression of essential spliceosome components.

### *C9ORF50* knockout impairs spliceosome function with increased aberrant alternative splicing events

To elucidate how C9ORF50 knockout inhibits spliceosome gene expression, we first evaluated chromatin accessibility using assay for transposase-accessible chromatin sequencing (ATAC-seq). The analysis revealed 606 sites with increased accessibility and 1204 sites with decreased accessibility in C9ORF50 knockout cells compared to NTC controls (**Figure 4A**). However, correlation analysis showed no clear segregation between C9ORF50 knockout and NTC cells (**Figure 4B**). Surprisingly, Gene Ontology (GO) analysis indicated that the most significantly accessible sites were enriched in RNA processing and splicing, RNP complex biogenesis, and spliceosomal snRNPs, suggesting increased transcriptional activity in these pathways (**Figure 4C**). For instance, key RNA processing genes Fus and Srsf5 exhibited increased chromatin accessibility in C9ORF50 knockout cells **(Figure S8**). Despite this, the reduced mRNA levels of these genes point to a compensatory mechanism, implying that additional post-transcriptional regulatory processes may contribute to the suppression of spliceosome gene expression.

**Figure 4.**
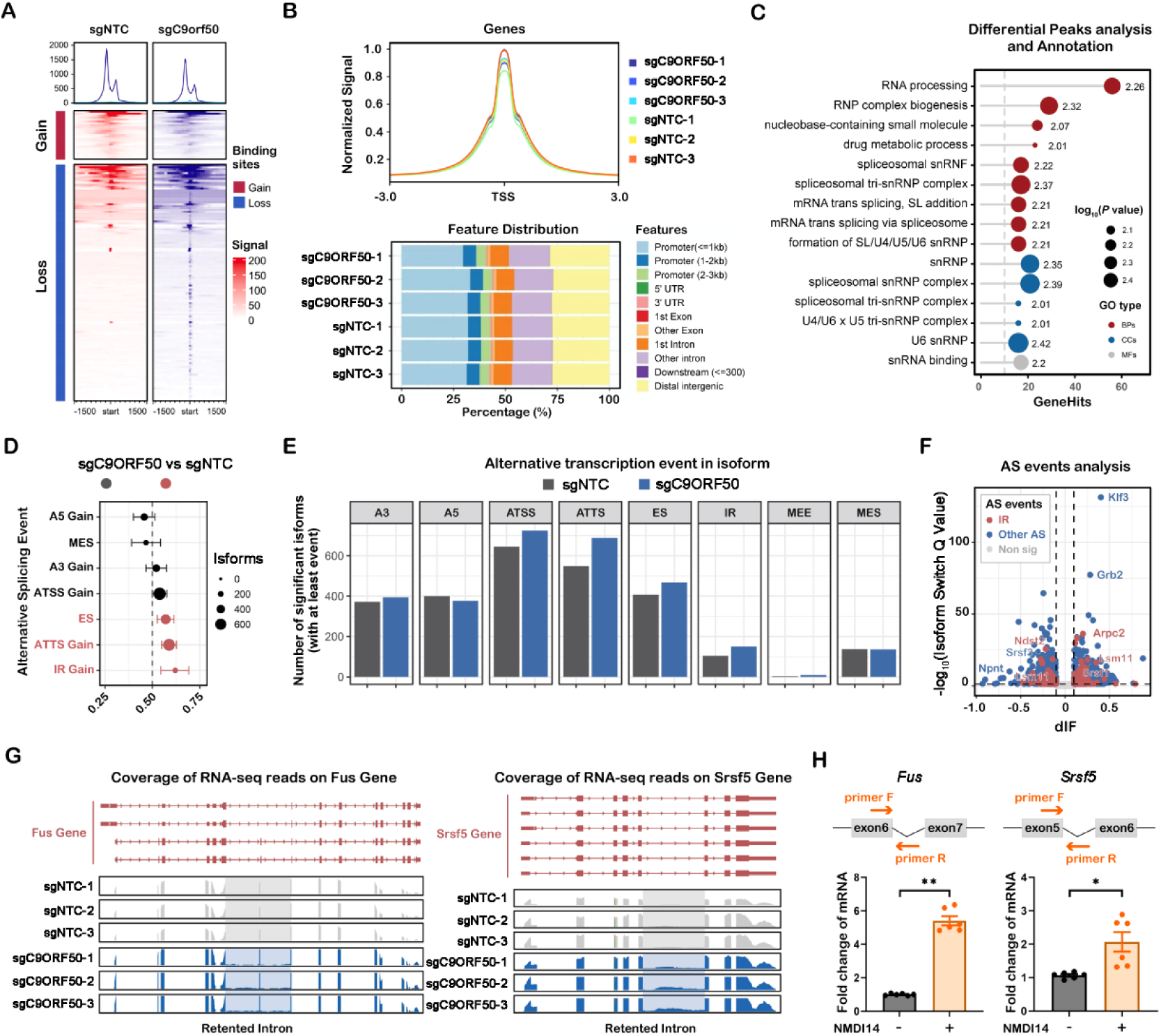
*C9ORF50* deficiency impairs spliceosome function via aberrant alternative splicing events in spliceosome-related genes. (A) Heatmap showing the ATAC-seq peaks classified on the basis of their changes in in MC38-sgNTC and MC38-sgC9ORF50 cells. Each row represents a locus (ATAC-seq peak center ± 1.5 kb), and the red gradient color indicates the ATAC-seq signal intensity. The numbers above the column indicate the replicates. (B) Metaplot showing the ATAC-seq signals at TSSs in MC38-sgNTC and MC38-sgC9ORF50 cells (upper). Annotation of the location of ATAC peaks in MC38-sgNTC or MC38-sgC9ORF50 samples to the reference genes (down). (C) GO pathway analysis of differential peak annotated genes. (D) Summary plot showing the enrichment of specific isoform features (consequence) resulting from the observed isoform switching events. The x-axis of the plot shows the fraction of switches having the indicated consequence, where <0.5 means depleted while >0.5 means enriched upon C9ORF50 knockout. (E) Number of detected alternative splicing events among the alternative splicing types. Intron retention (IR), exon skipping (ES), multiple exons skipping (MES), alternative transcription start sites (ATSS), alternative transcription termination sites (ATTS), alternative 5’ splice sites (A5), and alternative 3’ splice sites (A3). (F) Volcano plot showing the differential isoform fraction (dIF) and the significance of the switching isoforms. (G) Density of RNA-Seq reads (blue) mapping to exons (red) and introns in the *Fus* (J) and *Srsf5* (K) gene, showing retention of intron (marked by gray shadow). (H) The qPCR analysis the mRNA level of genes in C9ORF50 knockout MC38 cells after treated with/without NMDI14. The locations of the designed primers for *Fus* and S*rsf5* genes are shown: forward primer F; backward primer R. Boxes represent exon sequences, black lines represent intronic sequences. Data are shown as mean ± SEM, n = 3, Student’s t test (*p < 0.05, **p < 0.01).

Subsequently, we explored whether C9ORF50 knockout affects the processing of spliceosome-related transcripts. Using a differential transcript usage (DTU) analysis of RNA-seq data, we identified widespread aberrant alternative splicing events, including Intron retention (IR), exon skipping (ES), and alternative transcription termination sites (ATTS), in C9ORF50 knockout MC38 cells (**Figure 4D-4F**). In spliceosome-related genes like Fus and Srsf5, we observed an increased level of reads mapped to intronic regions and exon-intron boundaries, indicating defective splicing following C9ORF50 knockout (**Figure 4G**). IR can induce premature stop codons (PTCs), triggering nonsense-mediated mRNA decay (NMD)^29^. Indeed, treatment with the NMD inhibitor NMDI14 further elevated levels of intron-retaining spliceosome transcripts, suggesting these genes undergo NMD (**Figure 4H**). Moreover, RNA-seq results also showed that C9ORF50 knockout MC38 cells upregulated the expression of NMD genes (**Figure S7A**). These findings demonstrate that C9ORF50 knockout impairs the spliceosome’s ability to efficiently process pre-mRNA, resulting in reduced splicing efficiency and increased intron retention in the mRNA profiles.

### C9ORF50 knockout activates antitumor immunity via cytoplasmic accumulation of dsRNA

Given previous study linking spliceosome inhibition to the cytoplasmic accumulation of mis-spliced mRNAs forming double-stranded RNA (dsRNA) structures, which are recognized by dsRNA-binding proteins to activate apoptosis and antiviral pathways^30^, We investigated whether C9ORF50 knockout triggers similar responses. Immunofluorescence assays using the dsRNA-specific J2 antibody revealed a significant accumulation of cytoplasmic dsRNA in C9ORF50 knockout cells, which was abrogated by RNase III treatment, confirming the presence of dsRNA (**Figures 5A and 5B**). This dsRNA is likely detected by dsRNA recognition receptors, such as RIG-I and MDA5, initiating a signaling cascade that activates TANK binding kinase 1 (TBK1) and IκB kinase ε (IKKε), leading to the phosphorylation of interferon regulatory factor 3 (IRF3) and induction of type-I interferon (IFN) expression ^31^. Accordingly, western blot analysis confirmed increased phosphorylation of IRF3 and TBK1 following C9ORF50 knockout, indicative of their activation (**Figure 5C**). Moreover, key interferon-stimulated genes (ISGs), including Irf7, Isg15, Mx1, Oas1, as well as cytokines Ifnα, Ifnβ, Il-6, and Tnfα, were upregulated in C9ORF50 knockout MC38 cells compared to NTC controls (**Figure 5D-5F**). Enzyme-linked immunosorbent assay (ELISA) further confirmed the elevated secretion of IFNα, IFNβ, IL-6, and TNFα from C9ORF50 knockout cells (**Figure 5G**). Together, these findings indicate that C9ORF50 knockout activates the cytoplasmic dsRNA sensing pathway, leading to robust innate immune responses in cancer cells, potentially contributing to antitumor immunity.

**Figure 5.**
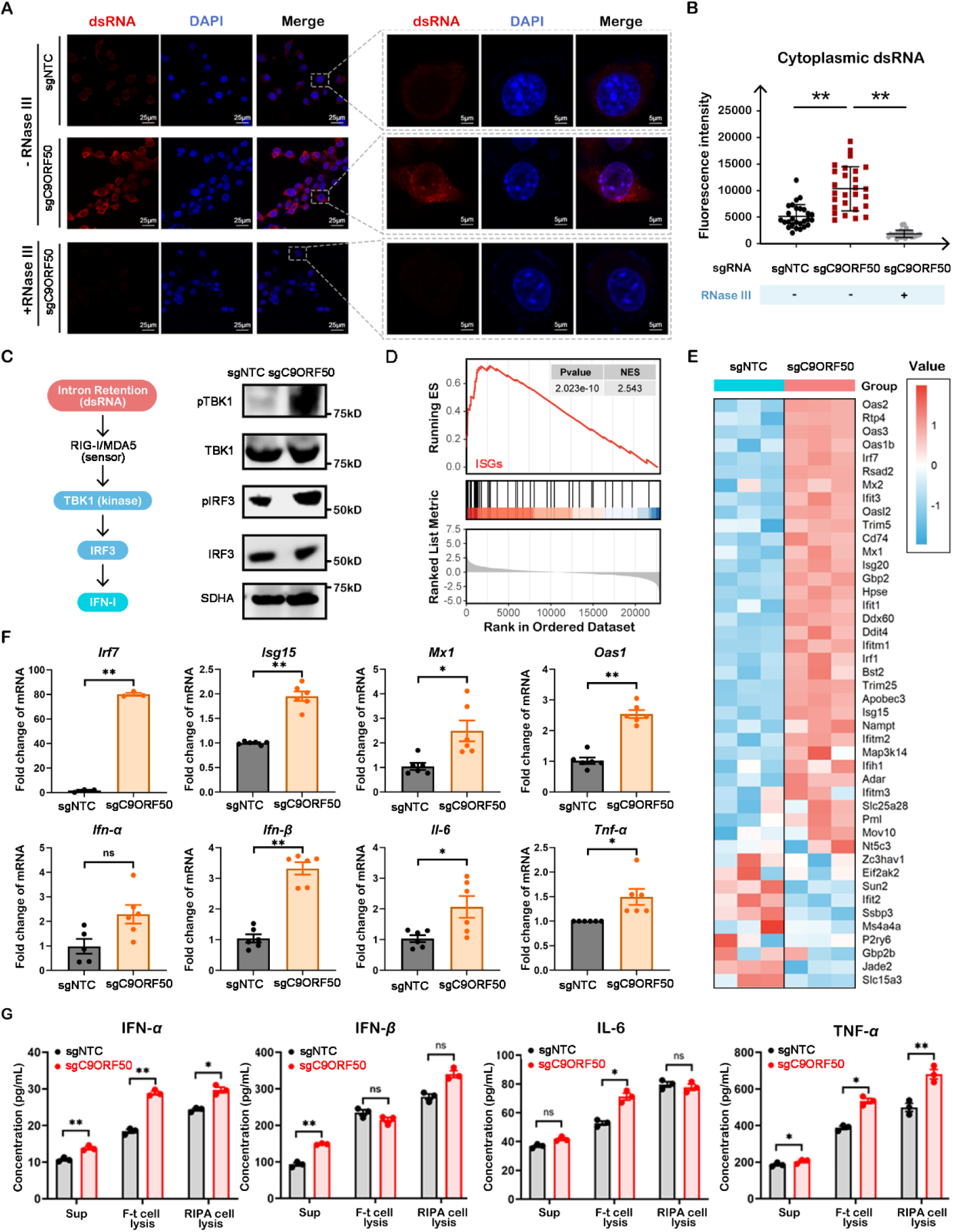
C9ORF50 knockout activates antitumor immunity through cytoplasmic accumulation of double-stranded RNA. (A) C9ORF50 deficiency induces cytoplasmic dsRNA accumulation. MC38-sgNTC and MC38-sgC9ORF50 cells were fixed and visualized by fluorescence microscopy using antibodies against *dsRNA* (red). RNase III treatment used as negative control for dsRNA signal. Scale bars, 25μm. Images representative of 3 experiments. (B) Quantification of cytoplasmic dsRNA signal intensity by ImageJ software. Data are shown as mean ± SEM, n = 34, Student’s t test (*p < 0.05, **p < 0.01). (C) C9ORF50 deficiency leads to increase of the phosphorylation of IRF3 and TBK1 in C9ORF50 knockout cells compared to in NTC cells. Protein levels of p-TBK1/TBK1, p-IRF3/IRF3, and SDHA were measured by Western blotting. SDHA served as loading control. (D) The enriched gene sets of ISG in C9ORF50 knockout MC38 cells compared with that in NTC MC38 cells are analyzed using GSEA. (E) Heatmap of the ISG in C9ORF50 knockout MC38 cells and NTC MC38 cells. (F and G) C9ORF50 deficiency leads to production of ISG. Gene expression was assayed by qPCR (F). Elisa validation experiments confirmed the expression in sgNTC-MC38 cells (G). Cell culture medium, cell lysis (RIPA-treated or freezing and thawing (F-t)) was collected, and the secretion levels of cytokines were detected by ELISA. Data are shown as mean ± SEM, n = 3, Student’s t test (*p < 0.05, **p < 0.01).

### *C9ORF50* knockout remodels the TME with increased T cell infiltration

To elucidate the impact of C9ORF50 knockout on the immune cell compositions of the TME, we performed an immune landscape profiling of syngeneic MC38 tumors with C9ORF50 knockout versus NTC controls. Flow cytometry analysis revealed a marked increase of CD45^+^ leukocyte infiltration in C9ORF50 knockout tumors (**Figure S9A**). Subsequent single-cell RNA sequencing (scRNA-seq) of the CD45^+^ tumor-infiltrating leukocytes, yielded a comprehensive transcriptional landscape of 39,544 leukocytes. Unsupervised clustering identified 18 distinct leukocyte clusters. Through deep learning-based and manual marker-based annotation, six major leukocyte subsets were identified: macrophages/monocytes, T cells, B cells, granulocytes, NK cells, and dendritic cells (**Figures 6A ans S9B**). Analyses of these subsets showed a significant increase in lymphoid infiltration in C9ORF50 knockout tumors (*p* = 0.033), particularly T cells (*p* = 0.081) and B cells (*p* = 0.048), along with a reduction in myeloid infiltration (*p* = 0.047), especially macrophages/monocytes (*p* = 0.025) (**Figure 6B**). By re-clustering of T cells, we characterized 11,775 cells across 13 distinct T cell subtypes (**Figure 6C**). C9ORF50 knockout led to significant increases in Th1 cells (P = 0.021), T follicular helper (Tfh) cells (P = 0.031), and CD8^+^ effector T cells (P = 0.025) compared to controls (**Figure 6D**), without significant changes in Treg cells.

**Figure 6.**
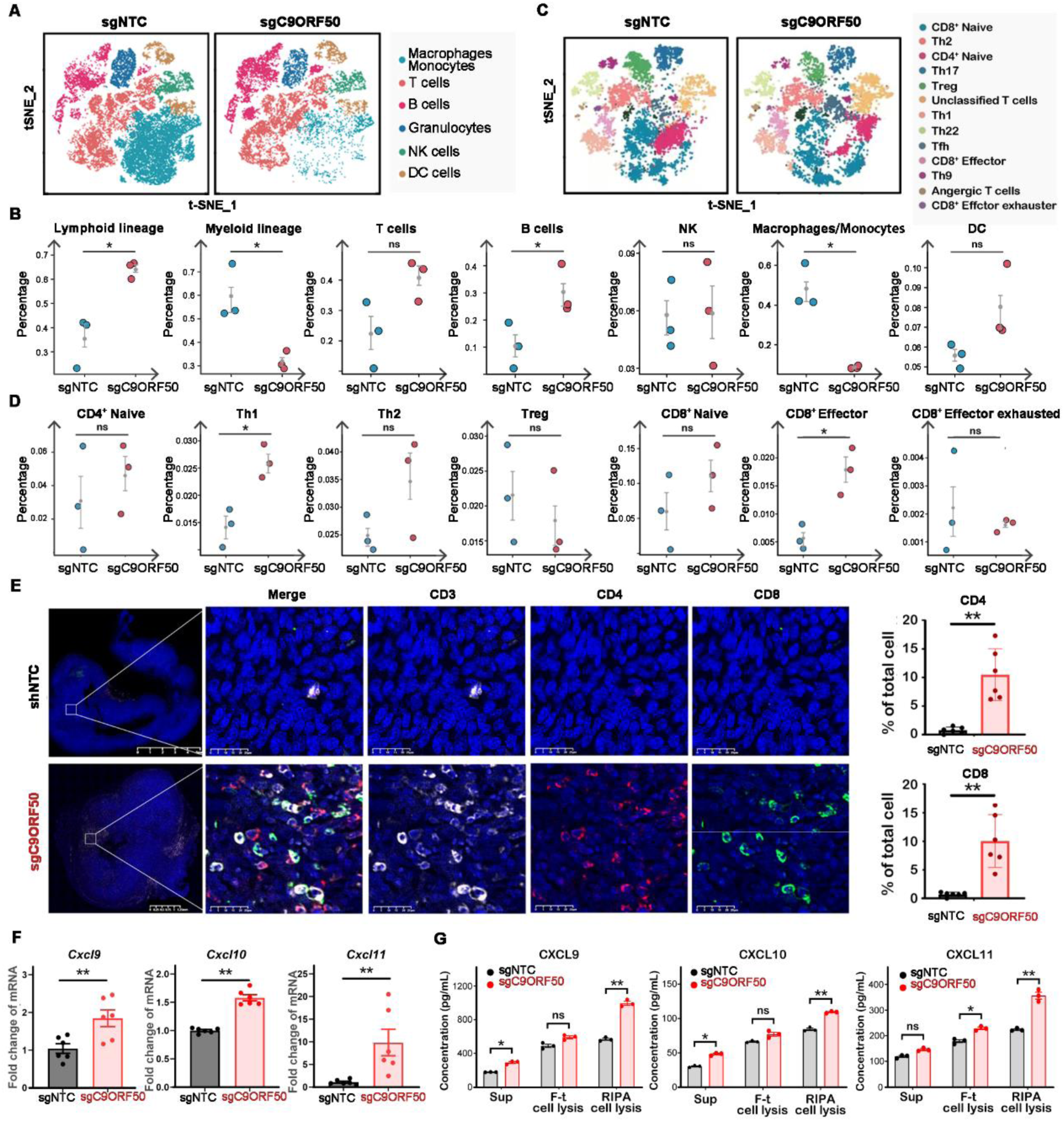
C9ORF50 knockout remodels the TME into an immune-stimulatory state. (A) Uniform manifold approximation and projection (UMAP) plot showing the clustering of different cell subsets in C9ORF50-deletion-driven tumor cells (sgC9ORF50, n=3 mice) or NTC tumor cells (sgNTC, n=3 mice). 6 major leukocyte cell types including Macrophages/Monocytes, T cells, B cells, Granulocytes, NK cells, and Dendritic cells are shown. (B) Comparing the differentiation potential of cell subpopulations in the sgNTC or sgC9ORF50 groups. The dots and box lines show subgroup scores. (C) Uniform manifold approximation and projection (UMAP) plot re-clustering and re-annotating T cells and other different cell subsets in the sgNTC or sgC9ORF50 groups. 13 major T cell types are shown. (D) Comparing the differentiation potential of cell subpopulations in the sgNTC or sgC9ORF50 groups. The dots and box lines show subgroup scores. (E) Representative CD3, CD4 and CD8 multicolor immunofluorescence staining in the sgNTC or sgC9ORF50 tumors. Quantification of CD4 and CD8 immunoreactivities. Scale bar, 25 μm. Images representative of 3 experiments. Data are shown as mean ± SEM, n = 6, Student’s t test (*p < 0.05, **p < 0.01). (F) C9ORF50 deficiency leads to production of chemokines. Gene expression was assayed by qPCR. Data are shown as mean ± SEM, n = 6, Student’s t test (*p < 0.05, **p < 0.01). (G) Elisa validation experiments confirmed the expression in sgNTC-MC38 cells. Cell culture medium, cell lysis (RIPA-treated or freezing and thawing (F-t)) was collected, and the secretion levels of chemokines were detected by ELISA. Data are shown as mean ± SEM, n = 3, Student’s t test (*p < 0.05, **p < 0.01).

Further differential expression analysis (DEA) in CD8^+^ T cells revealed an enrichment of genes involved in T cell receptor signaling pathway and immune response-regulation (**Figure S9C**). Similarly, DEGs in Th1 and Th2 cells were implicated in biogenesis of ribonucleoprotein complex and regulation of mRNA processing (**Figure S9D**). Moreover, macrophage infiltration, particularly the M2 subtype, was significantly reduced in C9ORF50 knockout tumors (**Figures S9E-S9G**). These results suggest that C9ORF50 knockout may reprogram the TME from an immune-refractory state to an immune-stimulatory state, characterized by enhanced infiltration of effector T cells and reduced immunosuppressive myeloid cells, especially M2 macrophages. This shift could potentially transform “cold” tumors into “hot” tumors, making them more susceptible to immune-mediated clearance. To validate these findings from scRNA-seq, we performed immunofluorescence assays in mouse syngeneic tumors. The results showed that significantly higher CD8^+^ and CD4^+^ T cell infiltration was observed in C9ORF50 knockout mouse tumors versus NTC control (**Figure 6E**).

Given the type-I ISGs include chemokines, such as CXCL9 and CXCL10, release for T cell recruitment ^32^, we hypothesized that C9ORF50 knockout enhanced T cell infiltration by altering chemokine production. Indeed, C9ORF50 knockout MC38 cells upregulated the expression of T cell chemo-attractants, specifically CXCL9, CXCL10, and CXCL11, as confirmed by RNA-seq, qPCR, and ELISA (**Figures 6F, 7G, S9H-J**). These findings indicate that C9ORF50 knockout or deficiency promotes T cell infiltration and activation by shaping the TME through elevated expression of T cell chemo-attractants.

To further investigate whether T cell immunity is pivotal for the anti-tumor effects of C9ORF50 knockout, we inoculated *C9ORF50* knockout and NTC control MC38 cells into T cell-deficient nude mice. C9ORF50 knockout only resulted in a modest reduction in tumor growth compared to controls (**Figure S10A**). Flow cytometric analysis revealed no significant differences in the overall infiltration of CD45^+^ immune cells, as well as the percentages of different innate immune cell subsets including neutrophils, monocytes, dendritic cells, macrophages and NK cells within CD45^+^ immune cells, between the two groups (**Figure S10B and S10C**). Further analysis demonstrated that C9ORF50 knockout does not significantly affect tumor cell proliferation (**Figures S11A and S11B**) but does substantially increase apoptosis, predominantly via activation of extrinsic apoptosis mediator caspase-3 and caspase-8 (**Figures S11C and S11D**), which may contribute to C9ORF50 knockout-resulted slightly reduced tumor growth in nude mice. Notably, this apoptotic mechanism aligns with prior findings that spliceosome inhibition could trigger extrinsic apoptosis via antiviral dsRNA-sensing pathways ^30^. Collectively, these findings suggest C9ORF50 deficiency primarily drives tumor regression by enhancing T cell infiltration, with a subsidiary contribution from extrinsic apoptosis in tumor cells.

### Tumor specific expression of C9ORF50 correlates with adverse clinical outcomes and immunosuppressive tumor microenvironment

To explore the clinical relevance of targeting C9ORF50, we analyzed its expression in human cancer cohorts and found that elevated C9ORF50 levels were inversely correlated with patient survival and significantly associated with advanced clinical staging in colorectal cancer, indicating its role in adverse clinical outcomes **(Figures 7A and 7B)**. Further analysis of 12 matched pairs of colorectal cancer and adjacent non-cancerous tissues revealed significant C9ORF50 upregulation in malignant tissues **(Figure 7C)**, while histological examination showed reduced cell density and increased leukocyte infiltration in C9ORF50^low^ tumors **(Figure 7D)**. Differential expression profiling via the ICRAFT platform^33^ highlighted a tissue-restricted expression signature of C9ORF50, with predominant abundance in malignant cells and minimal expression in immune cells and healthy tissues, underscoring its potential as a pharmacologically targetable molecule with limited off-target effects **(Figure 7E)**.

**Figure 7.**
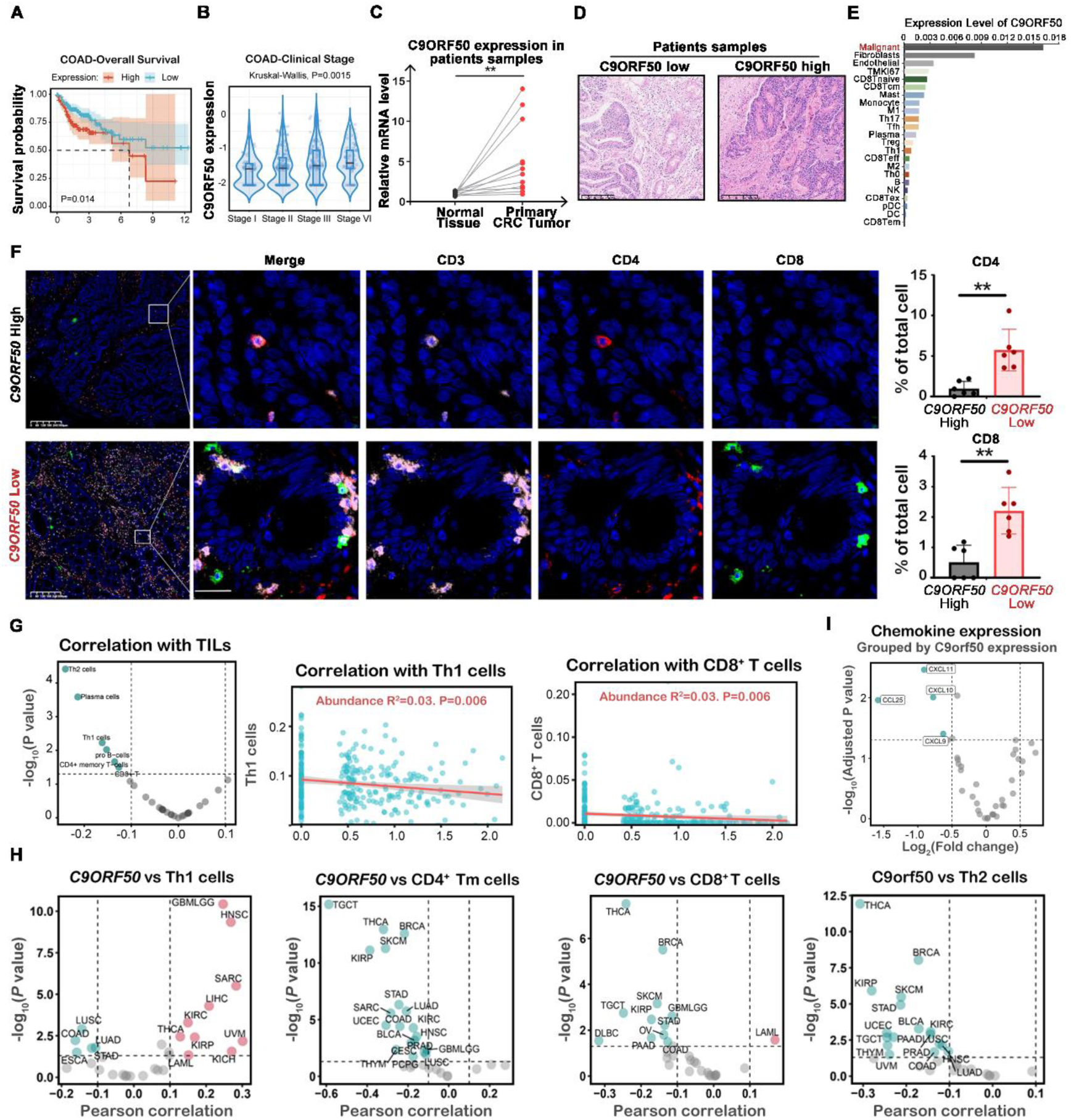
Tumor specific expression of C9ORF50 correlates with adverse clinical outcomes and immunosuppressive tumor microenvironment. (A) Kaplan–Meier overall survival analysis of colorectal cancer patients, based on the C9ORF50 expression. The p value was obtained by log-rank test. (B) Gene expression of C9ORF50 in colorectal cancer patients according to clinical stage. (C) C9ORF50 gene expression differences between 12 colorectal cancer patients’ samples. **P < 0.01 compared with normal samples. (D) Representative H&E images of tumors in (h). Scale bar, 100 μm. (E) The expression profile of C9ORF50 in malignant cells, immune cells, and healthy tissues was analyzed using the ICRAFT platform. C9ORF50 gene expression differences between 12 colorectal cancer patients’ samples. **P < 0.01 compared with normal samples. (F) Representative CD3, CD4 and CD8 multicolor immunofluorescence staining in C9ORF50 expressed high (upper) or low (down) human colorectal cancer tumors samples. Quantification of CD4 and CD8 immunoreactivities. Scale bar, 25 μm. Images representative of 3 experiments. The data represent the mean ± SEM, n = 3. (G) Volcano plot of the Spearman correlation between C9ORF50 mRNA expression and tumor infiltration immune cells (TILs), Th1, CD8^+^ T cells abundance in colorectal cancer. Green dots indicate immune cells significantly negatively correlated with C9ORF50 mRNA expression (adjusted P < 0.05). (H) Volcano plot of the Spearman correlation between C9ORF50 mRNA expression and Th1, Th2, CD8T and CD4 memory cells abundance in multiple cancer types. Red dots indicate immune cells significantly positively correlated with C9ORF50 mRNA expression (adjusted P < 0.05). Green dots indicate immune cells significantly negatively correlated with C9ORF50 mRNA expression (adjusted P < 0.05). (I) Volcano plot of the Spearman correlation between C9ORF50 mRNA expression and chemokines mRNA expression in colorectal cancer. Green dots indicate chemokines expression is significantly negatively correlated with C9ORF50 mRNA expression (adjusted P < 0.05).

To elucidate the potential role of C9ORF50 in shaping the tumor immune microenvironment, we performed immunofluorescence analysis of human colorectal cancer specimens. Our results demonstrated significantly enhanced infiltration of CD8^+^ and CD4^+^ T cells in C9ORF50^low^ tumor samples compared to their C9ORF50^high^ counterparts (**Figure 7F**). This observation was further corroborated by analysis of TCGA datasets, which revealed a consistent inverse correlation between C9ORF50 expression and infiltration of various lymphoid cell populations, including CD8^+^ T cells, Th1, Th2, CD4^+^ memory T cells, and plasma cells across multiple human cancer types particularly in colorectal cancer (**Figures 7G and 7H**). In addition, analysis of TCGA datasets demonstrated an inverse correlation between C9ORF50 expression and the transcriptional levels of key T cell chemoattractants (CXCL9, CXCL10, and CXCL11), consistent with our previously described mechanistic findings (**Figure 7I**). Collectively, these findings establish a compelling association between C9ORF50 overexpression and the establishment of an immunosuppressive TME, ultimately contributing to poor patient survival outcomes across multiple cancer types.

### Therapeutic targeting of C9ORF50 via RNA interference suppresses cancer progression

To evaluate the potential of C9ORF50 as a therapeutic target for cancer, we investigated the antitumor effects of C9ORF50 inhibition by direct intratumoral administration of *in vivo* siRNA in MC38 tumor-bearing mice (**Figure 8A**). First, we identified one siRNA with superior efficacy against C9ORF50 from three siRNAs (**Figure S12A**). Following intertumoral administration of a cholesterol-modified, *in vivo* stable C9ORF50 siRNA at a dose of 0.5 mg/kg, we observed a significant reduction in tumor burden in C9ORF50 siRNA treated mice compared to those from scramble siRNA control group (**Figure 8B**). Histopathological analysis showed reduced cell density and significant infiltration of inflammatory leukocytes in tumors treated with C9ORF50 siRNA (**Figure 8C**). Immunofluorescence staining further corroborated a marked increase in CD4^+^ and CD8^+^ T cell infiltration following C9ORF50 knockdown (**Figure 8D**). Interestingly, unmodified siRNA, although exhibiting a slightly less effective, also demonstrated notable antitumor effect (**Figures S12B-11F**). Taken together, these results suggested that C9ORF50 represents a novel and promising target for cancer therapy.

**Figure 8.**
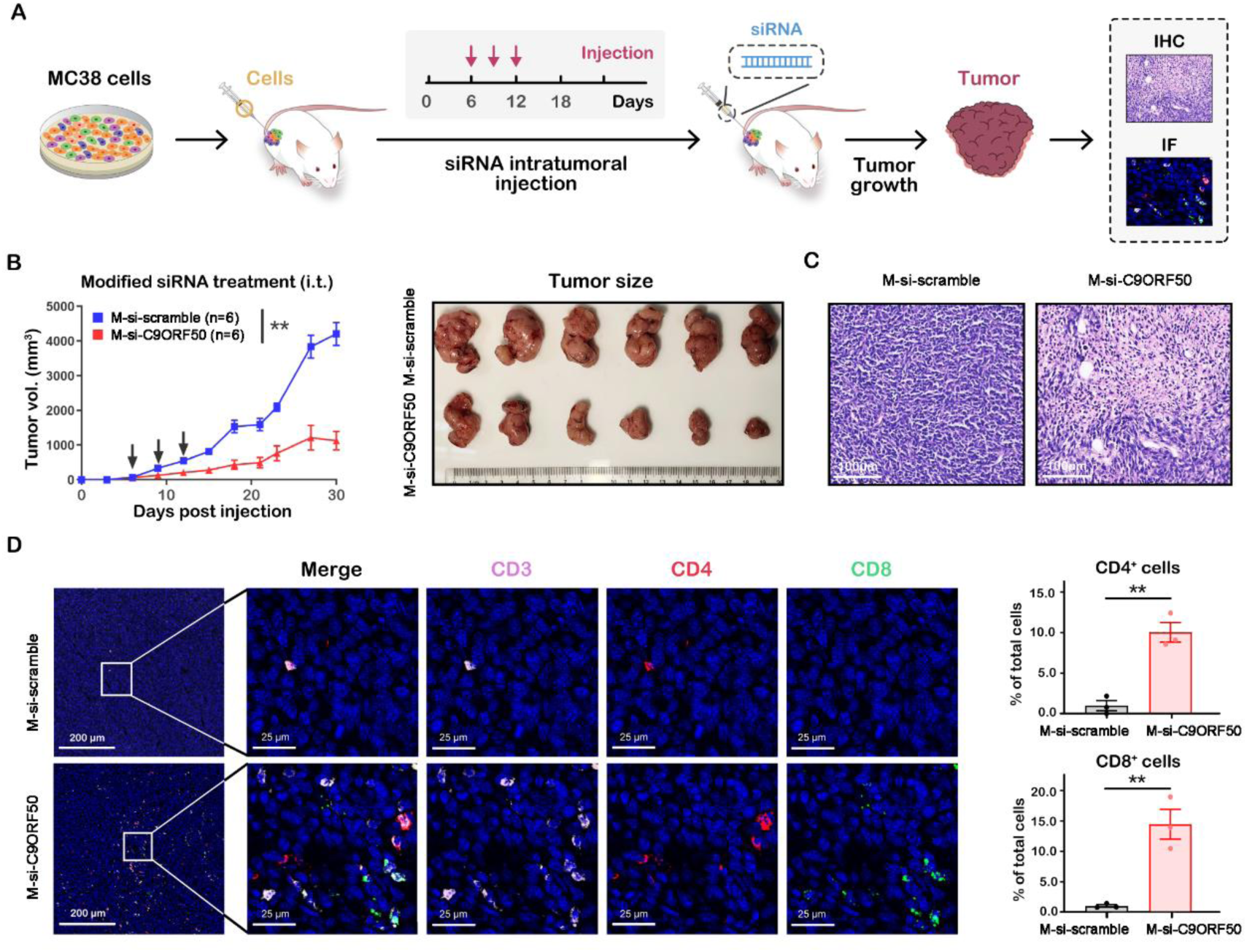
Therapeutic targeting of C9ORF50 via RNA interference suppresses cancer progression. (A) Schematic illustration of the immunotherapy mediated by *siRNA in vivo*. (B) Tumor growth curves after intratumoral injection (i.t.) of cholesterol-C9ORF50-siRNA (M-si-C9ORF50, n=6 mice) or cholesterol-scramble-siRNA (M-si-scramble, n=6 mice). **p < 0.01 by two-way ANOVA. Photographs of tumors from nude mice transplanted with C9ORF50-deletion-driven tumor cells. (C) H&E-stained tumor slices after different treatments. Scale bar, 25 μm. Images representative of 3 experiments. (D) Representative CD3, CD4 and CD8 multicolor immunofluorescence staining in M-si-C9ORF50 or M-si-scramble tumors. Scale bar, 25 μm. Images representative of 3 experiments. Quantification of CD3, CD4 and CD8 immunoreactivities. The data represent the mean ± SEM, n = 3. Student’s t test (**p < 0.01).

## Discussion

Despite the remarkable advancements in cancer immunotherapy, the majority of solid tumors remain refractory to these treatments, highlighting a critical need to address the underlying mechanisms of immune resistance. Cancer cell-intrinsic factors play a pivotal role in immune evasion by fostering an immune-privileged TME that excludes T cells and suppresses anti-tumor immune responses^4, 7, 9^. However, these factors remain largely unidentified, and uncovering the factors to counteract them is crucial for advancing cancer immunotherapy^7^. In this study, we conducted a genome-scale CRISPR knockout screen *in vivo* to identify potential factors to the cell-intrinsic immunity. This approach led to the identification of several intrinsic tumor factors, including well-established oncogenes such as Alk and Spi1 ^27, 28^, as well as several unrecognized factors. Notably, we identified C9ORF50, a previously uncharacterized gene, whose knockout resulted in significant tumor regression and improved survival in both mouse models and human patients. To our knowledge, no previous studies have explored the biological function of C9ORF50, while its aberrant DNA methylation has been proposed as early diagnostic biomarkers for colorectal and gastric cancers, implicating its involvement in gastrointestinal cancers ^34, 35^.

Our functional analyses reveal that C9ORF50 acts as a regulatory factor in spliceosome assembly by modulating LLPS and the expression of key spliceosome components. Since spliceosome inhibition has been shown to activate tumor immunity, targeting this complex is a promising cancer therapy strategy ^30^. However, the specific role of C9ORF50 in this process remains largely unexplored. In this study, we demonstrate that C9ORF50 deficiency or knockout leads to accumulation of intron-containing transcripts and aberrant dsRNA within the cytoplasm, triggering antiviral dsRNA-sensing pathways and extrinsic apoptosis. This disruption remodels the TME, dramatically increasing infiltration and activation of Th1 and CD8^+^ T cells, partially mediated by the upregulation of T cell chemo-attractants (**Figure S13**). Further analysis of clinical sample revealed a significant inverse correlation between C9ORF50 expression and both adaptive immune cell infiltration and patient prognosis, reinforcing its potential as a cancer therapeutic target. Notably, *in vivo* administration of a C9ORF50-targeting siRNA in syngeneic MC38 tumor model resulted in a dramatic reduction in tumor growth, underscoring C9ORF50 as a promising and novel target for cancer therapy.

Alternative splicing alterations are a common characteristic of cancer, affecting key tumor behaviors, such as proliferation, apoptosis, invasion, metastasis, and therapeutic response ^36, 37^. Small molecules targeting the spliceosome has gained prominence as a promising anti-cancer treatment. Previous research has demonstrated that the inhibition of U2 snRNP-associated spliceosomes components leads to alterations in splicing and gene expression of transcripts that are enriched for RNA-processing factors. This finding establishes a connection between fundamental splicing mechanisms and their effectors with modulators of alternative splicing and RNA processing ^38^. Based on this, we hypothesize that C9ORF50 inhibition or knockout disrupts phase separation within the spliceosome complex, impairing the proper pre-mRNA splicing, especially the splicing of essential spliceosome components like Fus, Srsf5, Srsf1, Sf3b1, Lsm6 and Hnrnpm. This feedback-enhanced disruption exacerbates spliceosome inhibition, creating a positive feedback loop that induces widespread alternative splicing dysregulation to activate antitumor immune responses.

A significant challenge in cancer therapy is minimizing off-target effects associated with systemically administered drugs, which often limit efficacy or cause unacceptable toxicity ^39^. To address this challenge, tissue-specific drug delivery systems or therapeutic targets with tissue-specific expression patterns need to be developed. Notably, C9ORF50 shows a highly tissue-specific expression, predominantly expressed in the gastrointestinal tract, brain, and kidneys, with minimal expression in immune cells, as well as immune cell-rich organs such as the spleen, thymus, liver, and lung ^40^. This distinct expression profile is consistent with our mouse tissue distribution analysis, underscoring C9ORF50’s potential role in tissue-specific biological processes. Furthermore, the correlation between C9ORF50 overexpression and poor patient prognosis further highlights its clinical relevance, though future studies should determine whether its oncogenic role extends across multiple cancer types through pan-cancer analyses.

In summary, our study has uncovered C9ORF50 as a previously recognized regulator of spliceosome assembly and as a novel target to activate cancer-intrinsic innate immunity for cancer therapy. Our findings highlight C9ORF50 as therapy target holds potentials to reprogram the TME and convert “cold” tumors into “hot” tumors, enhancing their susceptibility to immune-mediated clearance. Moreover, this study also paves the way for further exploration into additional regulators that could unlock cancer cell-intrinsic mechanisms, thereby enhancing tumor immunogenicity. The conserved nature of the phase separation domain of C9ORF50 across vertebrate species suggests the necessity of exploring its involvement in other pathologies related to splicing. Further investigation is warranted to assess the potential compensatory effects of paralogs, such as members of the C9ORF50 family, as well as the long-term implications of chronic disturbances in spliceosome function. Moreover, the development of humanized models that incorporate patient-derived xenografts is essential to more accurately replicate the human immune environment.

## Acknowledgements

This study was supported by grants from the National Natural Science Foundation of China (32171429 (L.Z.), 31902401(T.S.)), the Natural Science Foundation of Hunan Province (2022JJ30672 (L.Z.), 2024JJ9218 (L.Z.)).

## Author contributions

T. S. and L. Z. designed the research; T. S., C. L., J. K., S. X, Y. Q., L. Z., F. L.,Y. L., M. L., Y. L., T.H., S. Z., J. W., Y. L., Y. Q., J. L., Y. H. performed the research; T. S., C. L., S. Z., L.Z., J. Z, A. F. and M. L. analyzed the data; T. S., G.W. and L. Z. wrote the manuscript.

## Methods

### Animals

Six- to eight-week-old C57BL/6 and nude mice in both sexes were sourced from commercial vendors and used as transplant hosts. All animals were maintained in standard individually ventilated, pathogen-free conditions, with a light–dark cycle of 12 h and temperature around 21–25 °C, humidity of 40–60%. All animals’ experiments follow ARRIVE guidelines and followed strict randomization.

### Cell lines

All cell lines, including HEK293T cells, MC38 (colorectal cancer cells), Hepa1-6 (Hepatoma cells), Pan02 (pancreatic cancer cells), E0771 (triple-negative breast cancer cells), and B16F10 (melanoma cells), were obtained from the National Infrastructure of Cell Line Resource (Beijing, China) and authenticated by STR profiling. All cells were cultured in Dulbecco’s Modified Eagle Medium (DMEM) or Roswell Park Memorial Institute (RPMI) 1640 supplemented with 10% Fetal Bovine Serum (FBS), 1% Penicillin-Streptomycin (Penstrep), and maintained in a humidified CO_2_ incubator set to 5% CO_2_ at 37°C. Regular testing using a Mycoplasma Detection Kit (Thermo Fisher Scientific) or conventional PCR ensured that all cell lines were free of Mycoplasma contamination. To mitigate the risk of phenotypic drift, all cell lines were cryopreserved in multiple aliquots upon receipt.

### Genetic CRISPR knockout

Target gene knockout cells were generated using lentivirus-mediated CRISPR-Cas9 technology. sgRNAs specific for target genes including *C9ORF50*, as well as non-targeting control sgRNAs (sgNTC), were designed using the online CRISPR design tool (F. Zhang lab, MIT, Boston, MA). The sgRNA sequences were cloned into plentiCRISPR-v2 using the standard protocol. To generate the knockout cell lines, tumor cells were initially transduced with a lentiviral Cas9 to establish Cas9-stably expressing cell lines. These Cas9 stably expressing cell lines were then transduced with lentiviruses carrying either the sgC9ORF50 or sgNTC expressing sequences, generating *C9ORF50*-knockout cells and NTC cells, respectively.

### Genome-wide CRISPR-Cas9 knockout screening in vivo

The genome-wide mBrie CRISPR-knockout (KO) pooled library was purchased from Addgene (#73632) and amplified according to the supplier’s protocol. Cas9-expressing MC38 (MC38-Cas9) cells were transduced with the lentiviruses carrying mBrie library at a multiplicity of infection (MOI) of 0.2, ensuring 500x coverage of the library. Selection of transduced cells commenced 24 hours post transduction, utilizing 1.0 μg/ml puromycin for a period of 7 days, culminating in the establishment of a comprehensive genome knockout library cell line, designated MC38-mBrie. Following one week of *in vitro* cultivation, both the MC38-mBrie cells and the NTC cells were inoculated subcutaneously into the right rear groin of C57BL/6 mice at a density of 16 × 10^6^ cells per site. Tumor growth in the mice was closely monitored, with measurements taken every 3 days using vernier calipers. On day 10 post-inoculation, the mice were humanely euthanized, and the tumors were harvested for analysis. Simultaneously, an equivalent number of MC38-mBrie cells were cultured *in vitro* for 10 days, serving as a control group for the screening process. Genomic DNA from all samples was extracted using the Universal Genomic DNA Purification Mini Spin Kit (Qiagen catalog #56304). The sgRNA libraries were read out through a two-step PCR strategy. In the initial PCR (PCR#1), a region encompassing the sgRNA cassette was amplified with primers detailed in Supplementary Table 1, ensuring sufficient genomic DNA was included to maintain the full library complexity. The second PCR (PCR#2) utilized the pooled PCR#1 products, employing barcoded secondary primers (as listed in Supplementary Table 1) to incorporate the necessary sequencing adapters. The PCR#2 products were pooled and normalized for each biological replicate before merging the uniquely barcoded samples. The combined product was purified using the QiaQuick kit (QIAGEN) and quantified using a Nanodrop spectrometer. The libraries were subsequently diluted and sequenced on the NovaSeq 6000 platform (Illumina).

### Demultiplexing and Read Preprocessing

Raw single-end fastq read files were filtered and demultiplexed using Cutadapt. The following settings: cutadapt –discard-untrimmed -a GTTTTAGAGCTAGAAATGGC was used to remove extra sequences downstream of the sgRNA spacer sequences. As the forward PCR primers used to readout sgRNA representation were designed to have a variety of barcodes to facilitate multiplexed sequencing, we then demultiplexed these filtered reads with the following settings: cutadapt -g file: fbc.fasta–no-trim, where fbc.fasta contained the possible barcode sequences within the forward primers. Finally, the following settings: cutadapt –discard-untrimmed –g GTGGAAAGGACGAAACACCG was used to remove extra sequences upstream of the sgRNA spacers. Through this procedure, the raw fastq read files could be pared down to the 20 bp sgRNA spacer sequences.

### Mapping of spacers and quantitation of sgRNAs

Having extracted the 20 bp sgRNA spacer sequences from each demulitplexed sample, we then mapped the sgRNA spacers to the mBire library (Supplementary Table1). To do so, we first generated a bowtie index of either sgRNA library using the bowtie-build command in Bowtie1.1.2. Using these bowtie indexes, we mapped the filtered fastq read files using the following settings: bowtie-v 1–suppress 4,5,6,7–chunkmbs 2000 – best. Using the resultant mapping output, we quantitated the number of reads that had mapped to each sgRNA within the library. To generate sgRNA representation bar plots, we set a detection threshold of 1 read, and counted the number of unique sgRNAs present in each sample.

### Normalization and Summary-Level Analysis of sgRNA Abundances

The number of reads in each sample was normalized by converting raw sgRNA counts to reads per million (rpm). The rpm values were then subject to log_2_ transformation for certain analyses. To generate correlation heatmaps, we used the NMF R package and calculated the Pearson correlations between individual samples using log_2_ rpm counts. We first averaged the normalized sgRNA counts across all samples within a given group to calculate the cumulative distribution function for each sample group. To generate empirical cumulative distribution plots, we then used the ecdfplot function in the latticeExtra R package.

### RIGER and MAGeCK analyses

The enrichment of sgRNA and the corresponding gene were calculated with RNAi Gene Enrichment Ranking (RIGER) and the Model-based Analysis of Genome-wide CRISPR/Cas9 Knockout (MAGeCK) as described^41^. For RIGER analysis, read count tables were used to calculate log fold changes for tumor versus cell samples in order to score and rank sgRNAs, with ties in rank broken by random order. This data was then used as input to a Java-based implementation of RIGER (https://github.com/broadinstitute/rigerj) in order to generate p values and gene rankings based on consistent enrichment across multiple sgRNAs for identification of candidate genes. Both the second highest-ranking sgRNA and the weighted sum scoring methods were used for computation of gene rankings. For MAGeCK analysis, read count tables were used as inputs to a command-line-based tool (https://sourceforge.net/p/mageck/wiki/Home/) with the treatment group defined as the tumor samples and the control group defined as the cell pellet samples, with a list of non-targeting control sgRNAs provided for normalization and generation of the null distribution of RRA. Native MAGeCK plotting functions were used for visualization of RRA score and p value distributions and individual sgRNA read counts of selected genes. To generate Venn diagrams of overlapping enriched sgRNAs or genes, we considered all sgRNAs that were found to be significantly across RIGER and MAGeCK analyses.

### Orthotopic mouse model

Male C57BL/6 mice, 6-8 weeks old, were anesthetized, and a small incision was made to expose the intestine. Under microscopic visualization, a 50 µl suspension containing 2 × 10^6^ cells was injected into the intestine wall. The intestine was then returned to the peritoneal cavity, and the incision was closed with sutures. Mice were monitored for recovery and provided with appropriate analgesics. They were observed for any signs of distress or complications. Tumor growth was assessed at the end of the study period, typically 9 days post-implantation, by euthanizing the mice and performing necropsies to examine tumor progression.

### Subcellular localization and immunofluorescence assays

For subcellular localization assays, a transient expression vector, pCMV-C9ORF50-GFP was constructed. MC38 cells transiently transfected with pCMV-C9ORF50-GFP, were grown on glass coverslips. After 24h, cells were fixed with 4% formaldehyde in PBS for 10 min at 4 °C, followed by wash with PBS for 3 times. Coverslips were mounted on slides using anti-fade mounting medium with DAPI. Cells were visualized with a Leica TCS SP8 imaging system in fluorescence imaging mode. 405 Diode laser was used for DAPI detection; 488 Argon was used for GFP detection.

For cell immunofluorescence, pCMV-C9ORF50-GFP transfected MC38 cells were cultured in chamber slides overnight, and fixed with 4% formaldehyde in PBS for 10 min at 4 °C, followed by permeabilization with 0.5% Triton X-100 in PBS for 10 min. Cells were then blocked for nonspecific binding with 10% goat serum in PBS and 0.1% Tween-20 (PBST) overnight, and incubated with the anti-SF3B1 antibody (Abcam) or unrelated IgG for 1h at room temperature, followed by incubation with Alexa Fluor 568-labeled secondary antibody (Invitrogen) for 30 min at room temperature. Coverslips were mounted on slides using anti-fade mounting medium with DAPI. Immunofluorescence images were acquired on a Leica TCS SP8 imaging system in fluorescence imaging mode. 405 Diode laser was used for DAPI detection; 488 Argon was used for GFP detection; DPSS 561 was used for Alexa Fluor 568 detection.

### *In vitro* phase separation assay

The *in vitro* phase separation assay was conducted in storage buffer with specified protein concentrations, and PEG8000 was added to reach a final concentration of 10% (w/v). The phase separation assay was conducted on glass-bottom dishes (NEST), sealed with a clear adhesive film to prevent evaporation, and observed using an OLYMPUS FV3000 confocal microscope with 60× oil immersion lenses. The phase separation assay using human or mouse C9ORF50 was performed in a physiological LLPS buffer (20mM Tris-HCl, pH 7.5, 15mM NaCl, 130mM KCl, 5mM KH2PO4, 1.5mM MgCl2, and 1 mg/mL BSA).

### Fluorescence recovery after photobleaching

FRAP experiments were performed on an OLYMPUS FV3000 confocal microscope with a 60× oil immersion objective at 37 °C in a live-cell-imaging chamber. During the *in vitro* tests, droplets underwent photobleaching at 50% laser power for 0.5 seconds with 488-nm lasers. Time-lapse images were taken every three seconds for five minutes post-bleaching. For the *in vivo* experiments, droplets were bleached with a 488-nm laser pulse (50% intensity, 0.5 s). The recovery from photobleaching was recorded for the indicated time. Analysis of the recovery curves was carried out with FIJI/ImageJ software.

### Co-immunoprecipitation assay and mass spectrometry

HEK293T cells were transfected with 10 μg of either the GFP-tagged *C9ORF50* expression vector or a GFP-tag control plasmid. Protein harvesting occurred 48 hours post-transfection, with extraction carried out using RIPA lysis buffer supplemented with a protease inhibitor cocktail (Roche). After a 10-minute incubation on ice, the lysates were analyzed for protein concentration via the BCA method. Co-immunoprecipitation was performed using the Pierce Classic Magnetic IP/Co-IP kit (Thermofisher) according to manufacturer’s instructions. A total of 2 mg of protein was subjected to immunoprecipitation with 10 μg of rabbit anti-GFP antibody. The immunoprecipitated complexes were then separated on SDS-PAGE followed by Coomassie Blue staining. Gel fragments were excised, detained in 50% ethanol and 5% acetic acid, dehydrated in acetonitrile, dried in a Speed vacuum, and digested with trypsin. Tryptic peptides were separated on a C18 column and analyzed by LTQ-Orbitrap Velos (Thermo). Proteins were identified using the search engine of the NCBI against the Human NCBI RefSeq protein databases. The peptides were extracted from the polyacrylamide and subjected to liquid chromatography-mass spectrometry (MS) analysis. The interacted proteins identified by mass spectrometry were further validated through co-immunoprecipitation and Western blotting.

### Western Blot

MC38-sgC9ORF50 and MC38-sgNTC cells were lysed with the Cell Lysis Buffer (Cell Signaling). After incubation on ice for a minimum of 10 minutes, the lysates were analyzed for protein concentration using the bicinchoninic acid (BCA) assay. For Western blot analysis, 20 μg of each protein sample was loaded and separated via SDS-PAGE, then transferred onto PVDF membranes (Merck Chemicals). Following membrane blocking, primary antibodies were applied overnight at the indicated concentrations in a 2% BSA-TBST solution: anti-Phospho-IRF3 (Ser396), anti-IRF3, anti-Phospho-TBK1 (Ser172), anti-TBK1, anti-SDHA, anti-GFP, anti-U2SURP (all from ThermoFisher), anti-SRSF12 (from Aifang biological). Subsequently, the membranes were incubated with secondary antibodies, goat anti-rabbit HRP and rabbit anti-mouse HRP (both from ThermoFisher), for 1 hour. Detection was performed using the SuperSignal™ West Pico PLUS chemiluminescent substrate (ThermoFisher).

### RNA-seq analysis

Total RNA was extracted from MC38-sgC9ORF50 and MC38-sgNTC cells using TRIzol reagent (Invitrogen), following the manufacturer’s instructions. Thereafter, the NEBNext Ultra RNA Library Prep Kit for Illumina (New England Biolabs) was employed to construct the sequencing libraries. Sample clustering was performed using the TruSeq PE Cluster Kit v3-cBot-HS (Illumina), and sequencing was conducted on the Illumina NovaSeq 6000 platform. Differential expression analysis was conducted on three biological replicates per group, utilizing the DESeq2 R package (version 1.16.1). DESeq2 provided statistical routines for determining differential expression in digital gene expression data, employing a model based on the negative binomial distribution. The resulting *P* value was adjusted using the Benjamini and Hochberg approach for controlling the false discovery rate. Genes with an adjusted P < 0.05, as determined by DESeq2, were assigned as differentially expressed. Gene function enrichment analysis was conducted with clusterProfiler by R.

### Quantitative PCR assay

Total RNA was extracted from sgC9ORF50 and sgNTC MC38 cells using TRIzol reagent (Invitrogen) and subsequently reverse transcribed into complementary DNA (cDNA) with SuperScript III Reverse Transcriptase (Invitrogen), following the manufacturer’s protocol. Quantitative real-time PCR (qPCR) was conducted in 96-well plates using the SYBR Green PCR Master Mix (Roche). The fluorescence intensity was measured using a Light Cycler 480 instrument (Roche). Prime sequences for amplification are detailed in Supplementary Table 1.

### Detection and quantitation of cytokines and chemokines

MC38-sgC9ORF50 and MC38-sgNTC cells were cultured in standard media for 48 hours. Supernatants and whole-cell extracts were collected, with the latter obtained either through the freeze-thaw lysis method or using RIPA buffer. To prevent protein degradation, a protease inhibitor cocktail was incorporated into all samples. The concentrations of cytokines including IFNα, IFNβ, IL-6 and TNFα, as well as chemokines including CXCL9, CXCL10 and CXCL11, were quantified using specific enzyme-linked immunosorbent assay (ELISA) kits (Mouse IL-6 ELISA Kit, Mouse IFN-α ELISA Kit, Mouse IFN-β ELISA Kit, Mouse TNF-α ELISA Kit, Mouse CXCL9 ELISA Kit, Mouse CXCL10 ELISA Kit and Mouse CXCL11 ELISA Kit).

### ATAC-seq Analysis

ATAC-seq libraries were prepared following established protocols ^42^. Cell populations of MC38-sgC9ORF50 and MC38-sgNTC, each at a density of 1×10^4^ cells, were lysed, and accessible chromatin was tagged by transposition with the Nextera Tn5 transposase. The resulting transposed DNA fragments were purified using the Qiagen MinElute kit and subjected to amplification for 5-10 cycles with Nextera PCR primers. The ATAC-seq libraries were then submitted for paired-end sequencing on an Illumina NovaSeq 6000 platform. The raw FASTQ reads obtained were mapped to the mm10 genome assembly using Bowtie v2, and accessible genomic regions were identified with MACS2. These accessible regions across all samples were consolidated, and the read counts within each region were tallied. Pairwise Spearman correlation coefficients were calculated based on the read counts within each region to assess the reproducibility and similarity between samples. Differential accessibility analysis was conducted using DESeq2, which evaluates statistical significance in chromatin accessibility between conditions. The intersection of accessible regions with motif analysis was performed using HOMER, providing insights into the transcription factor binding events associated with these regions.

### Flow cytometry and cell sorting

Both sgC9ORF50 and sgNTC transduced MC38 cells were inoculated subcutaneously into the right flanks of C57BL/6 or nude mice, respectively, at a density of 2 × 10^6^ cells per site. Tumor tissues were harvested, minced, and treated with a digestion solution comprising PBS with 2% FBS, collagenase type IV (0.25 mg/mL, Sigma), and DNase I (20 U/mL, Sigma). The mixture was then agitated on a shaker incubator at 37 °C for 10 minutes to facilitate disaggregation, followed by vigorous pipetting with a 10-mL pipette. The resulting cell suspension was strained through a 70-μm nylon mesh to obtain a single-cell suspension. The single-cell suspensions were incubated with a rat anti-mouse CD16/CD32 monoclonal antibody (ThermoFisher) to block Fc receptor-mediated reactions, followed by the surface staining at 4 °C for 15 min. A panel of antibodies (BioLegend) was used for surface staining: APC-CD45, PE-CY7-CD11b, percp/cy5.5-MHC-II IA/IE, eFluor450-Ly6c, PE-F4/80, PE/Dazzle-CD11c, eFluor506-L/D and FITC-NKP46. The samples were analyzed using a fluorescent-activated cell sorter (FACS) Aria III cell sorter (BD Biosciences) with BD FACSDiva Software (v9.0, BD Biosciences). Data analysis was conducted using FlowJo software (Ashland).

### Cell apoptosis and proliferation analyses

Cell apoptosis was measured by annexin V-FITC/propidium iodide (PI) assay using flow cytometry as previously described. Specifically, cells were subjected to staining with the Annexin V-FITC/PI apoptosis detection kit (Yeasen) and subsequently analyzed using a FACS Aria III cell sorter (BD Biosciences). In the cell proliferation assay, 5-ethynyl-2’-deoxyuridine (EdU) was introduced to the culture medium at a final concentration of 1 μM, added 2 hours prior to detection and maintained until cell collection. At the designated time points, cells were trypsinized and processed for the Click-iT EdU assay following the manufacturer’s instructions (ThermoFisher). The proliferating cells were then quantified using the FACS Aria III cell sorter (BD Biosciences).

### Tissue H&E and immunofluorescence staining

For H&E staining, paraffin-embedded tumor tissue sections were deparaffinized in xylene and subjected to a graded ethanol series for rehydration. Antigen retrieval was performed using heat in citrate buffer. Subsequently, the tumor tissue sections were stained with hematoxylin for 1 minute, followed by a brief rinse and eosin staining for 1 minute. After additional rinsing, the slides were mounted with neutral balsam for preservation. For immunofluorescence, sections were also deparaffinized, rehydrated and subjected to antigen retrieval. The slides were then blocked with 5% goat serum in phosphate-buffered saline (PBS), and subsequently incubated with primary antibodies specific for CD3 (ThermoFisher), CD4 (ThermoFisher), and CD8 (ThermoFisher), alongside a rabbit IgG isotype control (ThermoFisher). Nuclei were counterstained with DAPI. A TSA indirect kit (PerkinElmer) was used according to the manufacturer’s instructions. Imaging was conducted using the Vectra® Polaris™ Imaging System (Akoya Biosciences), and subsequent image analysis was performed utilizing the HALO™ Image Analysis Software.

### scRNA-seq analysis

Tumor tissues from the MC38-NTC (n=3) and MC38-sgC9ORF50 (n=3) models were harvested from tumor-bearing mice to prepare single-cell suspensions. Tumor-infiltrating leukocytes were enriched through fluorescence-activated cell sorting (FACS), specifically isolating CD45^+^ cells to focus on immune cell populations. The cell suspensions were then counted and loaded onto a 10X Genomics Chromium platform for single-cell RNA sequencing, following the manufacturer’s protocol.

The analysis of the generated sequencing data was performed using Cell Ranger (version 4.0.0), which facilitated sample demultiplexing, barcode processing, alignment to the mouse genome (GRCm38), filtering of low-quality reads, counting of unique molecular identifiers (UMIs), and aggregation of sequencing runs. The resulting UMI count matrix was subsequently processed using Seurat (version 4.3.0) for downstream analyses. Low-quality cells were filtered based on the following criteria: a minimum of 200 detected genes and an upper limit of 7000 detected genes. Additionally, cells with more than 30% mitochondrial UMI counts or more than 0.3% hemoglobin UMI counts were excluded. Normalization and variance stabilization of the count data using the SCTransform function in Seurat. For data integration across different experimental groups, we employed the SCT integration method. The constructed shared nearest neighbor (SNN) graph was generated using the FindNeighbors function, and clustering was executed with the FindClusters function, setting the resolution parameter to 0.5. To visualize the clustered data further, we implemented UMAP and t-SNE for non-linear dimensionality reduction. UMAP was performed using the first 30 PCA dimensions, while t-SNE was executed on the same set of dimensions, with results plotted using the DimPlot function. The FindAllMarkers function in Seurat was utilized to identify differentially expressed features that characterize each cluster.

For cell annotation, we employed both database-based and marker-based approaches. The database-based auto-annotation utilized the SingleR package along with ImmGenData and MouseRNAseqData reference datasets. In the marker-based auto-annotation, we prepared a list of specific marker genes corresponding to various cell lineages, including lymphoid, myeloid, epithelial, and stromal cells. Leveraging the results from these different annotation methods, clusters were assigned to specific cell types based on the expression of key marker genes.

### RNA interference assay

Small interfering RNAs (siRNAs) targeting *C9ORF50*, as well as control scramble siRNAs, were designed and synthesized by GenePharma (Shanghai, China). To enhance intracellular stability, a cholesterol-modified version of the C9ORF50-siRNA (cholesterol-C9ORF50-siRNA) and a corresponding cholesterol-modified scramble siRNA were also produced. The efficacy and specificity of these siRNAs were rigorously validated using qPCR, and the siRNA demonstrating the optimal knockdown efficiency was selected for subsequent functional assays. The complete sequences are detailed in Supplementary Table 1.

Wild-type MC38 cells were injected subcutaneously into C57BL/6 mice. On the sixth day following tumor transplantation, mice were randomized into groups (n=5 per group) to receive either scramble siRNA, C9ORF50-siRNA, cholesterol-C9ORF50-siRNA, or cholesterol-scrambled siRNA by intratumoral injection. The injection formulation contained 30 μg of siRNA per mouse, administered every 2 days for a total of three injections. Throughout the treatment period, the tumor-bearing mice were closely observed, and tumor dimensions were measured every 3 days using vernier calipers in a blinded manner relative to cage labels. Following euthanasia, the tumors were harvested, weighed, and subjected to H&E staining as well as immunofluorescence analysis.

### Clinical samples

Colorectal cancer and paired normal tissue samples (located >5 cm from the malignant region) were obtained during surgery. All patients underwent resection of the primary tumor at the second Xiangya hospital of central south university (Changsha, China) and affiliated hospitals. Informed consent was obtained from all participants. We have complied with all relevant ethical regulations for work with human participants. The study protocols were approved by the ethics committee of central south university (Approval No. Z0449-01).

### Protein structure prediction and functional annotation

The sequence of C9ORF50 and its homologs were obtained from the UniProt database (https://www.uniprot.org/). Multiple sequence alignments were performed using the MUSCLE algorithm (https://toolkit.tuebingen.mpg.de/tools/muscle) and visualized with the R package ggmsa and Adobe Illustrator. The sequence of C9ORF50 was submitted to ColabFold (AlphaFold2) for structure prediction. Additionally, the AlphaFold3 web server (https://alphafoldserver.com/) was used for further structural analysis.

### Statistics and reproducibility

Data between two groups were analyzed using a two-tailed unpaired t-test. Multiple t-test using the Holm-Sidak method was used for multiple group comparison. Different levels of statistical significance were accessed based on specific *p* values and type I error cutoffs (0.05, 0.01, 0.001). GraphPad Prism Software and RStudio were used for these analyses.

### Code availability

Codes used for data analysis or generation of the figures related to this study are available from the corresponding author upon reasonable request.

## Supplementary Tables

Table S1 sheet 1. The sgRNA sequence of mBrie library

Table S1 sheet 2. Primers used in this work

## References

1. Dagher, O.K., Schwab, R.D., Brookens, S.K. & Posey, A.J. Advances in cancer immunotherapies. CELL 186, 1814 (2023).

2. Fan, T. et al. Therapeutic cancer vaccines: advancements, challenges, and prospects. SIGNAL TRANSDUCT TAR 8, 450 (2023).

3. Kyrysyuk, O. & Wucherpfennig, K.W. Designing Cancer Immunotherapies That Engage T Cells and NK Cells. ANNU REV IMMUNOL 41, 17–38 (2023).

4. Tang, L., Huang, Z., Mei, H. & Hu, Y. Immunotherapy in hematologic malignancies: achievements, challenges and future prospects. SIGNAL TRANSDUCT TAR 8, 306 (2023).

5. Sadeghi, R.H. et al. Understanding the tumor microenvironment for effective immunotherapy. MED RES REV 41, 1474–1498 (2021).

6. Ghorani, E., Swanton, C. & Quezada, S.A. Cancer cell-intrinsic mechanisms driving acquired immune tolerance. IMMUNITY 56, 2270–2295 (2023).

7. van Weverwijk, A. & de Visser, K.E. Mechanisms driving the immunoregulatory function of cancer cells. NAT REV CANCER 23, 193–215 (2023).

8. Wellenstein, M.D. & de Visser, K.E. Cancer-Cell-Intrinsic Mechanisms Shaping the Tumor Immune Landscape. IMMUNITY 48, 399–416 (2018).

9. Li, Y.R., Halladay, T. & Yang, L. Immune evasion in cell-based immunotherapy: unraveling challenges and novel strategies. J BIOMED SCI 31, 5 (2024).

10. Zhu, M. et al. Evasion of Innate Immunity Contributes to Small Cell Lung Cancer Progression and Metastasis. CANCER RES 81, 1813–1826 (2021).

11. Klimeck, L., Heisser, T., Hoffmeister, M. & Brenner, H. Colorectal cancer: A health and economic problem. BEST PRACT RES CL GA 66, 101839 (2023).

12. Cohen-Eliav, M. et al. The splicing factor SRSF6 is amplified and is an oncoprotein in lung and colon cancers. J PATHOL 229, 630–639 (2013).

13. Wan, L. et al. Splicing Factor SRSF1 Promotes Pancreatitis and KRASG12D-Mediated Pancreatic Cancer. CANCER DISCOV 13, 1678–1695 (2023).

14. Deng, L. et al. SF3A2 promotes progression and cisplatin resistance in triple-negative breast cancer via alternative splicing of MKRN1. SCI ADV 10, eadj4009 (2024).

15. Ciesla, M. et al. m(6)A-driven SF3B1 translation control steers splicing to direct genome integrity and leukemogenesis. MOL CELL 83, 1165–1179 (2023).

16. Danson, S.J. et al. Phase I pharmacokinetic and pharmacodynamic study of the bioreductive drug RH1. ANN ONCOL 22, 1653–1660 (2011).

17. Steensma, D.P. et al. Phase I First-in-Human Dose Escalation Study of the oral SF3B1 modulator H3B-8800 in myeloid neoplasms. LEUKEMIA 35, 3542–3550 (2021).

18. Corrionero, A., Minana, B. & Valcarcel, J. Reduced fidelity of branch point recognition and alternative splicing induced by the anti-tumor drug spliceostatin A. GENE DEV 25, 445–459 (2011).

19. Darrigrand, R. et al. Isoginkgetin derivative IP2 enhances the adaptive immune response against tumor antigens. COMMUN BIOL 4, 269 (2021).

20. Xu, Y., Nijhuis, A. & Keun, H.C. RNA-binding motif protein 39 (RBM39): An emerging cancer target. BRIT J PHARMACOL 179, 2795–2812 (2022).

21. Lu, S.X. et al. Pharmacologic modulation of RNA splicing enhances anti-tumor immunity. CELL 184, 4032–4047 (2021).

22. Reber, S. et al. The phase separation-dependent FUS interactome reveals nuclear and cytoplasmic function of liquid-liquid phase separation. NUCLEIC ACIDS RES 49, 7713–7731 (2021).

23. Peng, Q. et al. SRSF9 mediates oncogenic RNA splicing of SLC37A4 via liquid-liquid phase separation to promote oral cancer progression. J ADV RES (2025).

24. Calabretta, S. & Richard, S. Emerging Roles of Disordered Sequences in RNA-Binding Proteins. TRENDS BIOCHEM SCI 40, 662–672 (2015).

25. Gong, H., Xue, B., Ru, J., Pei, G. & Li, Y. Targeted Therapy for EWS-FLI1 in Ewing Sarcoma. CANCERS 15 (2023).

26. Moller, E. et al. EWSR1-ATF1 dependent 3D connectivity regulates oncogenic and differentiation programs in Clear Cell Sarcoma. NAT COMMUN 13, 2267 (2022).

27. Lou, L., Zhang, P., Piao, R. & Wang, Y. Salmonella Pathogenicity Island 1 (SPI-1) and Its Complex Regulatory Network. FRONT CELL INFECT MI 9, 270 (2019).

28. Shaw, A.T. & Engelman, J.A. ALK in lung cancer: past, present, and future. J CLIN ONCOL 31, 1105–1111 (2013).

29. Ge, Y. & Porse, B.T. The functional consequences of intron retention: alternative splicing coupled to NMD as a regulator of gene expression. BIOESSAYS 36, 236–243 (2014).

30. Bowling, E.A. et al. Spliceosome-targeted therapies trigger an antiviral immune response in triple-negative breast cancer. CELL 184, 384–403 (2021).

31. Chen, Y.G. & Hur, S. Cellular origins of dsRNA, their recognition and consequences. NAT REV MOL CELL BIO 23, 286–301 (2022).

32. Sistigu, A. et al. Cancer cell-autonomous contribution of type I interferon signaling to the efficacy of chemotherapy. NAT MED 20, 1301–1309 (2014).

33. Luo, C. et al. Integrated computational analysis identifies therapeutic targets with dual action in cancer cells and T cells. IMMUNITY 58, 745–765 (2025).

34. Cao, Y. et al. KCNQ5 and C9orf50 Methylation in Stool DNA for Early Detection of Colorectal Cancer. FRONT ONCOL 10, 621295 (2020).

35. Li, H. et al. Feasibility and reproducibility of a plasma-based multiplex DNA methylation assay for early detection of gastric cancer. PATHOL RES PRACT 238, 154086 (2022).

36. Bonnal, S.C., Lopez-Oreja, I. & Valcarcel, J. Roles and mechanisms of alternative splicing in cancer - implications for care. NAT REV CLIN ONCOL 17, 457–474 (2020).

37. Climente-Gonzalez, H., Porta-Pardo, E., Godzik, A. & Eyras, E. The Functional Impact of Alternative Splicing in Cancer. CELL REP 20, 2215–2226 (2017).

38. De Maio, A. et al. RBM17 Interacts with U2SURP and CHERP to Regulate Expression and Splicing of RNA-Processing Proteins. CELL REP 25, 726–736 (2018).

39. Zhao, Z., Ukidve, A., Kim, J. & Mitragotri, S. Targeting Strategies for Tissue-Specific Drug Delivery. CELL 181, 151–167 (2020).

40. Duff, M.O. et al. Genome-wide identification of zero nucleotide recursive splicing in Drosophila. NATURE 521, 376–379 (2015).

41. Li, W. et al. MAGeCK enables robust identification of essential genes from genome-scale CRISPR/Cas9 knockout screens. GENOME BIOL 15, 554 (2014).

42. Grandi, F.C., Modi, H., Kampman, L. & Corces, M.R. Chromatin accessibility profiling by ATAC-seq. NAT PROTOC 17, 1518–1552 (2022).

